## Supplemental Figures for "An Evolutionarily Conserved Regulatory Pathway of Muscle Mitochondrial Network Organization"

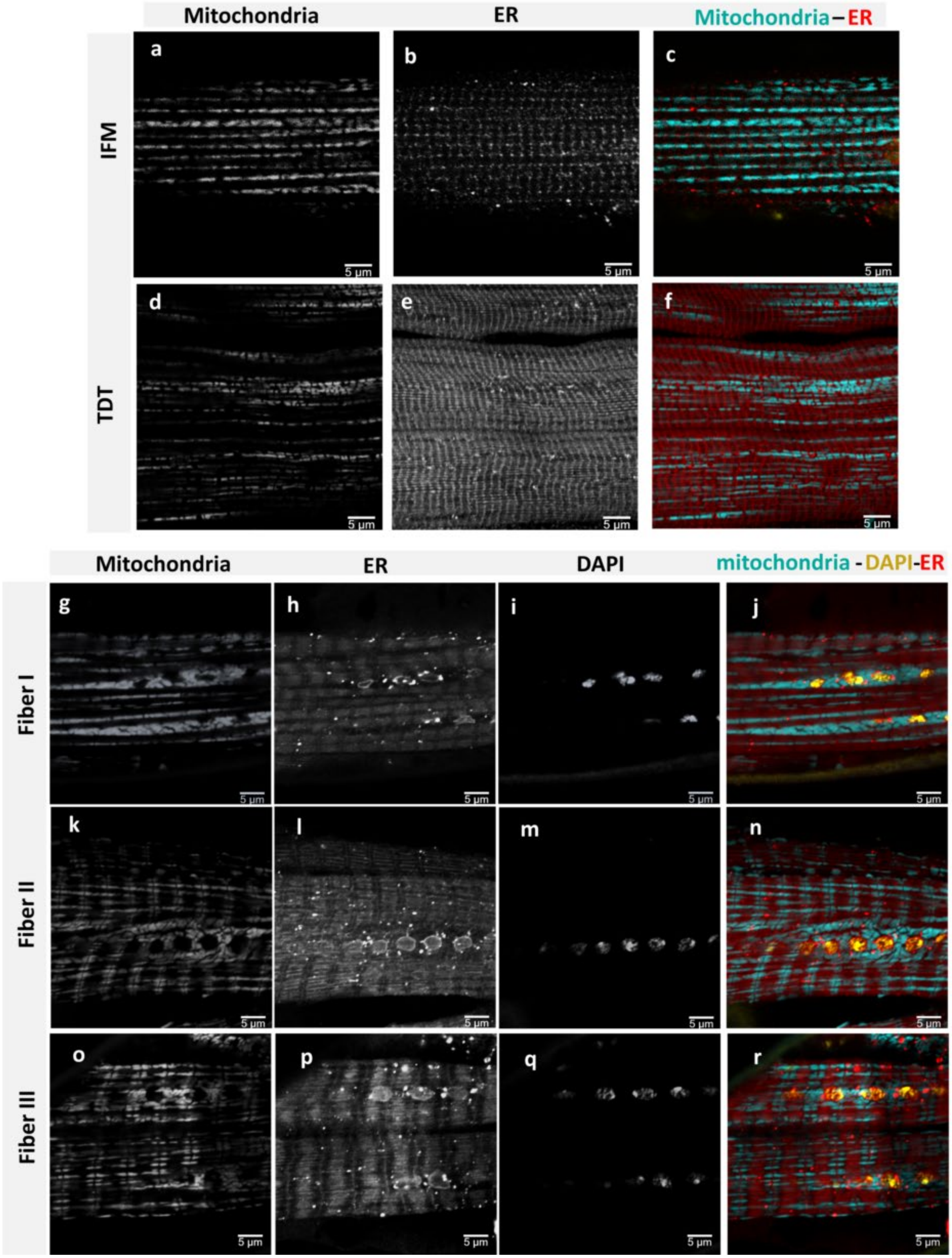


**Supplementary Fig. S1. Endoplasmic Reticulum in adult Drosophila muscles.** (a, b, c) Wildtype flight muscles (IFMs) showing thin endoplasmic reticulum (ER) (KDEL-rfp) and parallel mitochondria (mito-gfp). (d, e, f) Wildtype jump muscles (TDT) show abundant ER (KDEL-rfp) and parallel mitochondria (mito-gfp). (g-j) Wildtype Leg muscle Fiber I, (k-n) Fiber II, and (o-r) Fiber III stained for mitochondria (mito-gfp), ER (KDEL-rfp), and nuclei (DAPI) (Scale bars: 5 µm).

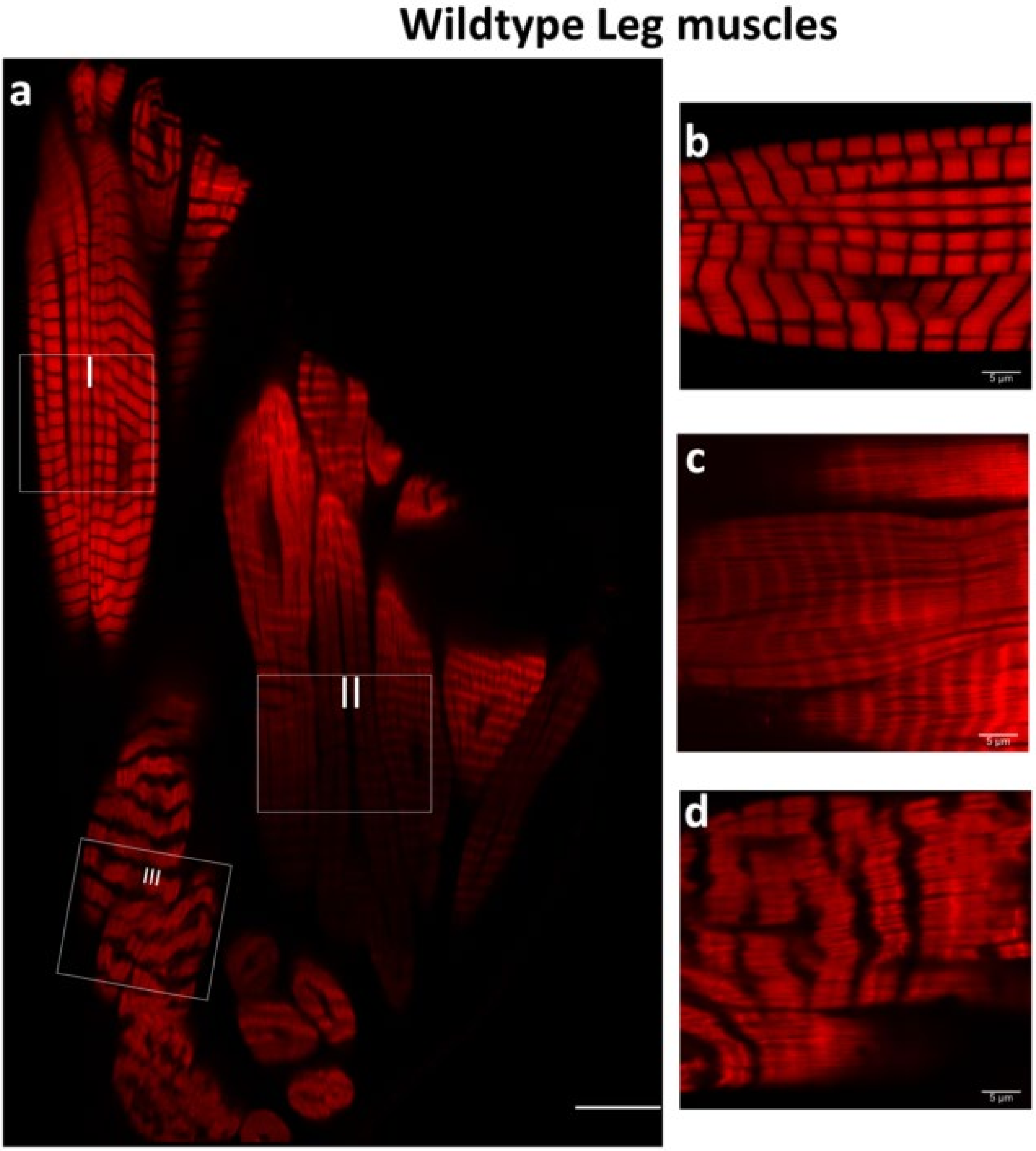


**Supplementary Fig. S2. Muscle organization in adult Drosophila leg muscles.** (a) Wildtype leg muscles (coxa) stained for F-actin (phTRITC) showing three distinct tubular muscle Fiber types (highlighted by squares, scale bar: 20 µm). (b) Tubular muscle fiber of Fiber I. (c) Tubular muscle fiber of Fiber II (d) Tubular muscle fiber of Fiber III (Scale bars: 5 µm).

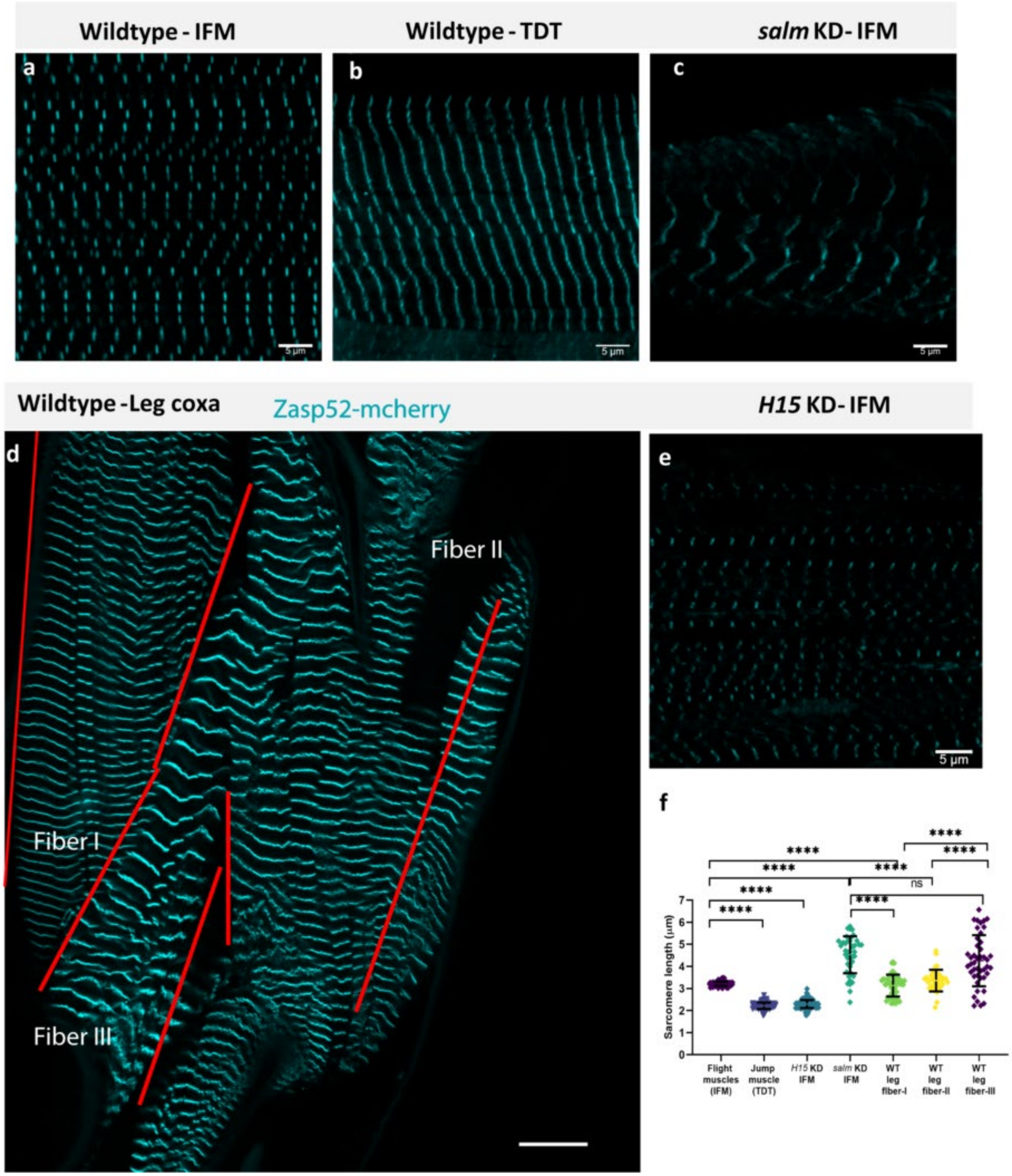


**Supplementary Fig. S3. Sarcomere organization in different muscles types in Drosophila.** (a, b, c) Sarcomeres labeled with zasp52-mcherry in (a) wildtype flight muscles (IFM), (b) wildtype jump muscles, and (c) *salm* KD IFM (Scale bars: 5 µm). (d) Wildtype leg muscles of coxa with sarcomeres labeled with zasp52-mcherry (Scale bar: 20 µm). (e) Sarcomeres labeled with zasp52-mcherry in *H15* KD IFM (Scale bar: 5 µm). (f) Quantification of individual sarcomere length in muscles (WT-IFM, *n=140*; WT-TDT, *n=139*; *H15* KD-IFM, *n=190*; *salm* KD-IFM, *n=43*; WT-Leg Fiber I, *n=56*; WT-Leg Fiber II, *n=37*; WT-Leg Fiber III, *n=44*). Each point represents individual sarcomere distance. Bars represent overall mean ± SD. Significance determined as $p < 0.05$ from one way ANOVA with Tukey's (*, $p \leq 0.05$; **, $p \leq 0.01$; ***, $p \leq 0.001$; ****, $p \leq 0.0001$; ns, non-significant).

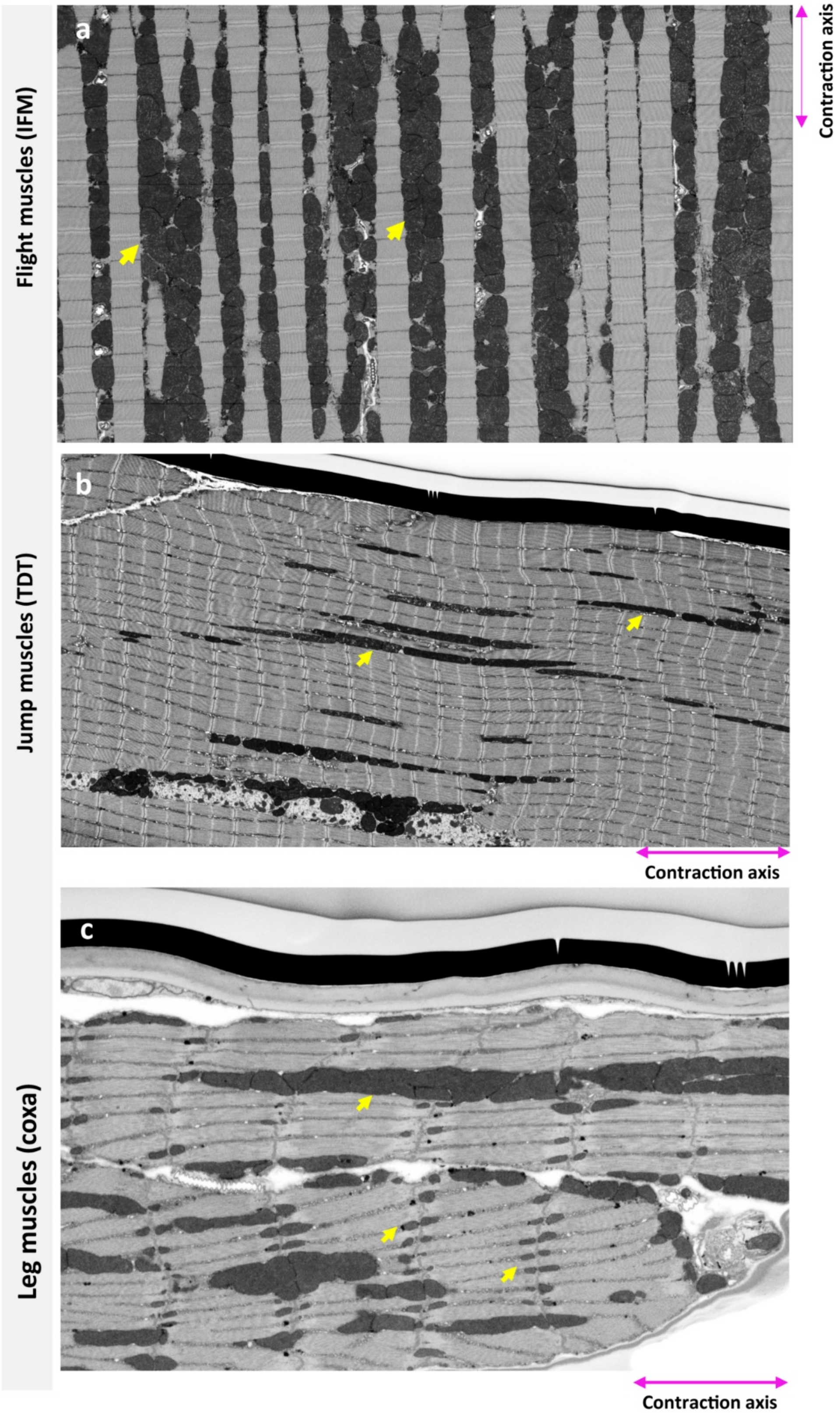


**Supplementary Fig. S4. Mitochondrial organization in adult Drosophila muscles.** (a, b, c) Electron micrographs showing mitochondrial organization in wildtype (a) flight, (b) jump, and (c) leg muscles. Arrows indicate mitochondria.

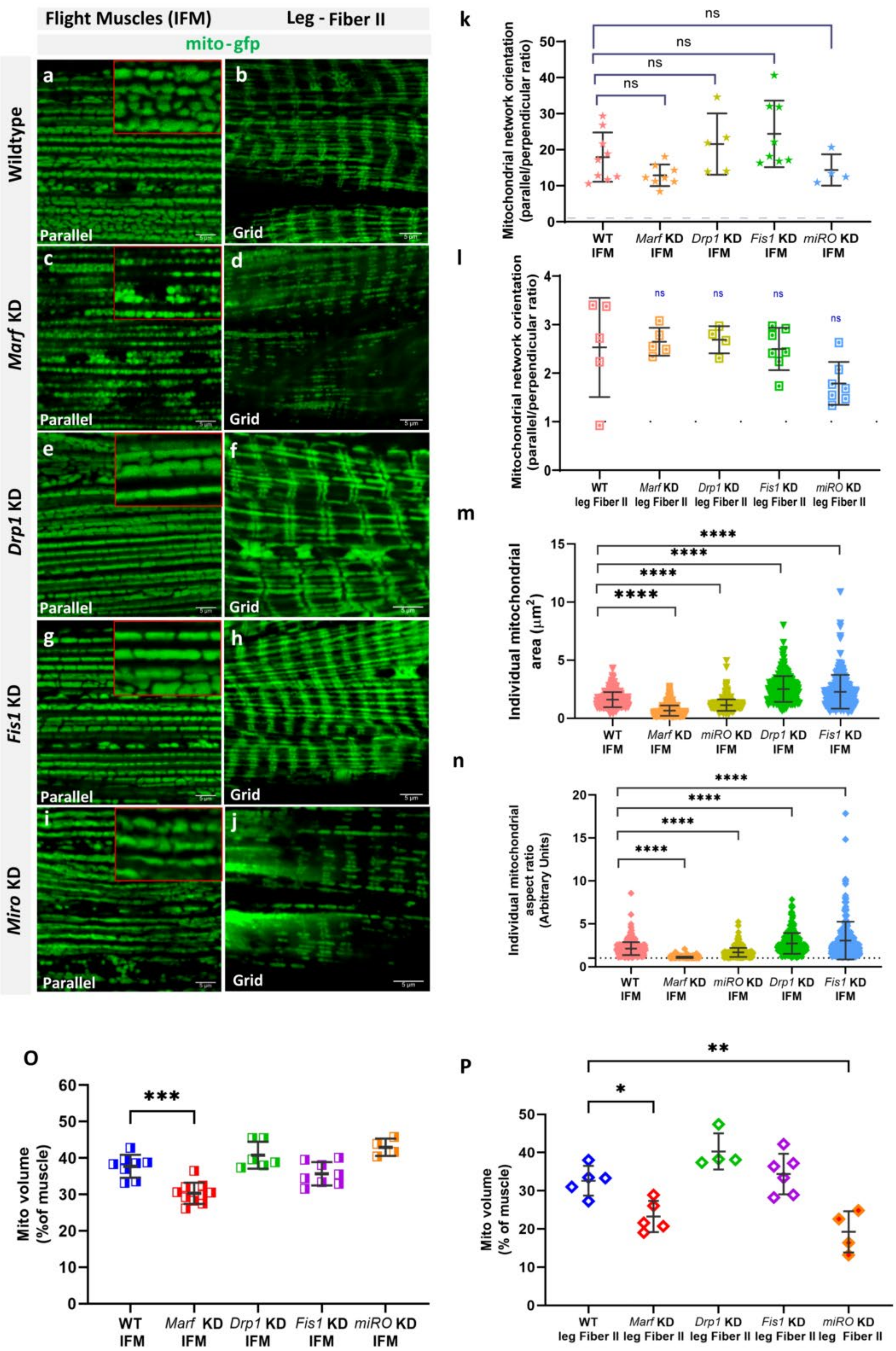

Flight Muscles (IFM)
Leg - Fiber II
mito-gfp
Wildtype
Marf KD
Drp1 KD
Fis1 KD
Miro KD
Parallel
Grid
k
Mitochondrial network orientation (parallel/perpendicular ratio)
ns
WT IFM
Marf KD IFM
Drp1 KD IFM
Fis1 KD IFM
miRO KD IFM
l
WT leg Fiber II
Marf KD leg Fiber II
Drp1 KD leg Fiber II
Fis1 KD leg Fiber II
miRO KD leg Fiber II
m
Individual mitochondrial area (μm²)
****
n
Individual mitochondrial aspect ratio (Arbitrary Units)
O
Mito volume (%of muscle)
***
P
Mito volume (% of muscle)
**
*

**Supplementary Fig. S5. Knockdown of mitochondrial dynamics regulators in Drosophila muscles does not affect mitochondrial networks.** (a) Adult wildtype flight muscles (IFM) stained for mitochondria (mito-GFP) reveal tubular mitochondria (inset image) and parallel mitochondrial networks. (b) Wildtype leg muscles showing grid mitochondrial networks (mito-GFP). (c) *Marf* KD in IFM results in circular and small individual mitochondria (inset image) that remain in parallel mitochondrial networks. (d) *Marf* KD in leg muscles shows grid-like mitochondrial networks. (e) *Drp1* KD in flight muscles shows longer mitochondria (inset image), but mitochondrial networks remain parallel. (f) *Drp1* KD leg muscles show grid-like mitochondrial networks. (g) *Fis1* KD in flight muscle shows parallel mitochondrial networks. (h) *Fis1* KD in tubular leg muscles shows grid-like mitochondrial networks. (i) Motility factor, *Miro* KD flight muscles shows abnormal mitochondria morphology (inset image), but retains parallel mitochondrial networks. (j) *Miro* KD in leg muscles showing grid-like mitochondrial networks (Scale bars: 5 µm). (k, l) Quantification of mitochondrial network orientation in (k) IFM and (l) leg muscle fibers. Dotted line represents parallel equal to perpendicular. *mito-gfp;mito-mcherry;Mef2-Gal4* used as Wildtype. (WT-IFM, *n=9*; *Marf* KD-IFM, *n=8*; *Drp1* KD-IFM, *n=5*; *Fis1* KD-IFM, *n=8*; Miro KD-IFM, *n=4*; WT-Leg Fiber II, *n=5*; *Marf* KD-Leg Fiber II, *n=5*; *Drp1* KD-Leg Fiber II, *n=4*; *Fis1* KD-Leg Fiber II, *n=7*; *Miro* KD-Leg Fiber II, *n=7*). Each point represents value for each dataset. (m) Quantification of individual mitochondrial area and (n) quantification of individual mitochondrial aspect ratio (major axis/minor axis) (WT-IFM, *n=393; Marf* KD-IFM, *n=1091*; *Miro* KD-IFM, *n=509; Drp1* KD-IFM, *n=329*; *Fis1* KD-IFM, *n=356*). Each point represents value for individual mitochondria. (o, p) Quantification of mitochondrial volume as a percent of total muscle volume in (o) IFM and (p) leg Fiber II. *UAS-mito-gfp;UAS-mito-OMM-mcherry;Dmef2-Gal4* used as wildtype. (WT-IFM, *n=8*; *Marf* KD-IFM, *n=9*; *Drp1* KD-IFM, *n=6*; *Fis1* KD-IFM, *n=8*; *Miro* KD-IFM, *n=4*; WT-Fiber II, *n=5*; *Marf* KD-Fiber II, *n=5*; *Drp1* KD-Fiber II, *n=4*; *Fis1* KD-Fiber II, *n=6*; *Miro* KD-Fiber II, *n=4*). Bars represent mean ± SD. Significance determined as $p < 0.05$ from one way ANOVA with Tukey's (*, $p \leq 0.05$; **,$p \leq 0.01$; ***,$p \leq 0.001$; ****, $p \leq 0.0001$; ns, non-significant).

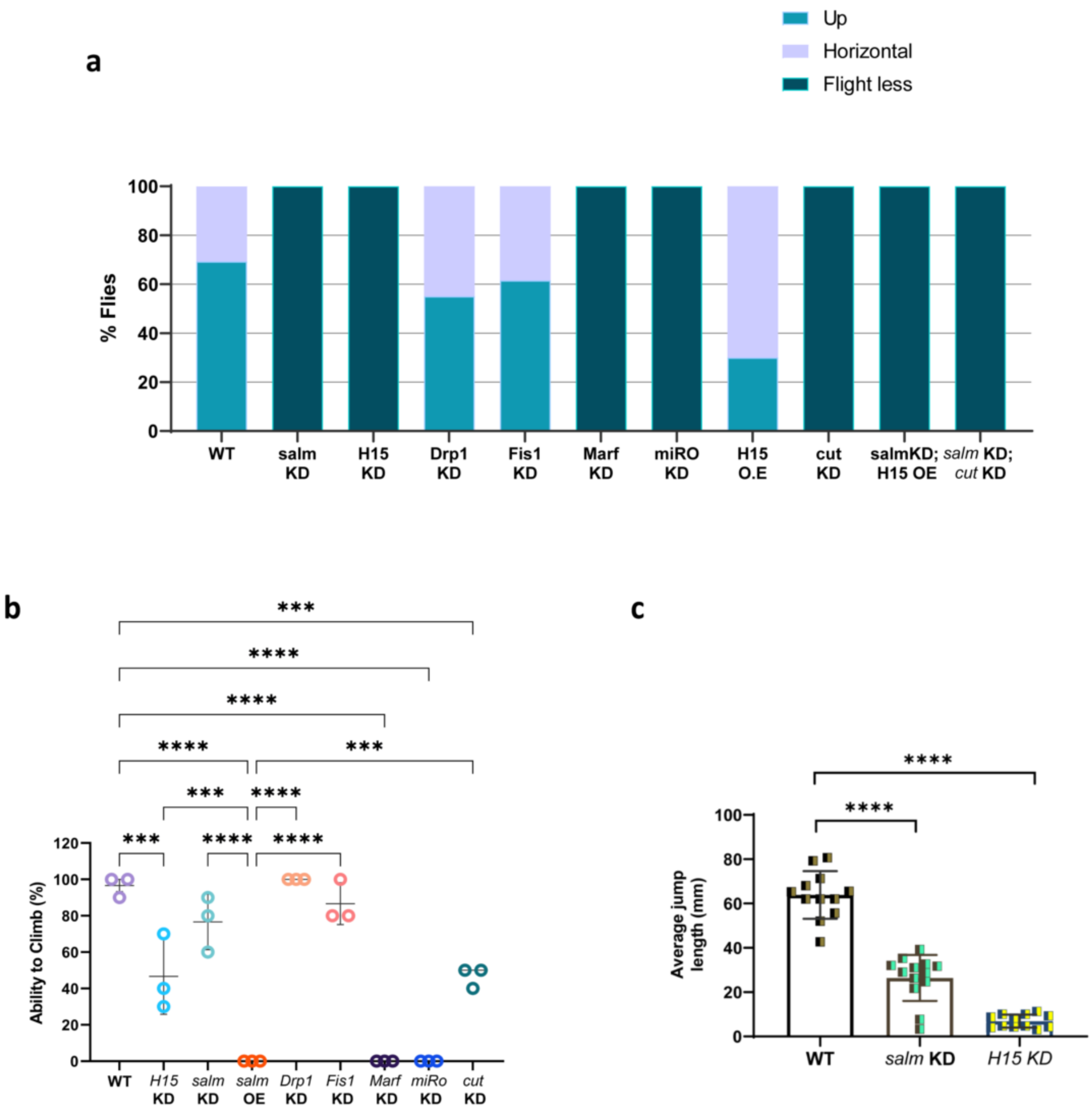


**Supplementary Fig. S6. Flight, climbing, and jump ability.** (a) Flight assay. *mito-gfp;Dmef2-Gal4* used as wildtype (WT, *n=13*; *salm* KD, *n=26*; *H15* KD, *n=22*; *Marf* KD, *n=25*; *Drp1* KD, *n=20*; *Fis1* KD, *n=13*; *Miro* KD, *n=13*; *H15* OE, *n=10; cut* KD, *n=20*, *salm* KD;*H15* OE, *n=13*; *salm* KD; *cut* KD, *n=24*). (f) Climbing assay. Each point represents one group (*n=10* for each group). (g) Average jump length (WT, *n=12*; *salm* KD, *n=13*; *H15* KD, *n=12*). Each point represents individual fly. Bars represent mean ± SD. Significance determined as $p < 0.05$ from one way ANOVA with Tukey's (*, $p \leq 0.05$; **,$p \leq 0.01$; ***,$p \leq 0.001$; ****, $p \leq 0.0001$; ns, non-significant).

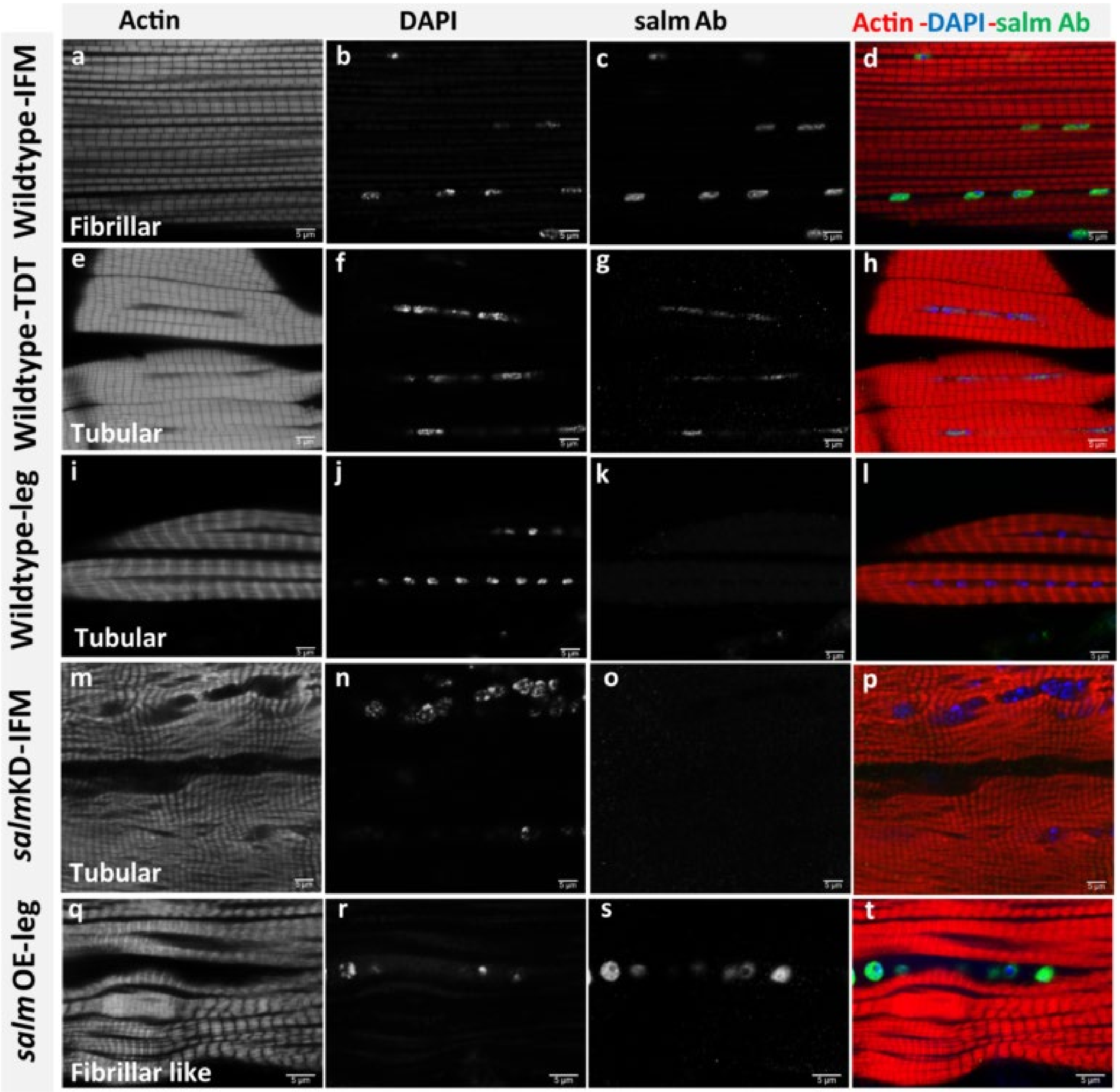


**Supplementary Fig. S7. Salm expression in Drosophila muscles.** (a-d) Wildtype flight muscles (IFM), (e-h) jump muscles, and (i-l) leg muscles stained for F-actin (phTRITC), nuclei (DAPI), and Salm antibody. (m-p) *salm* KD IFM showing decreased expression of Salm in nuclei (DAPI). (q-t) Leg muscles with *salm* OE stained for F-actin (phTRITC), nuclei (DAPI), and Salm antibody showing increased Salm expression in the nuclei (Scale Bars: 5 µm for all).

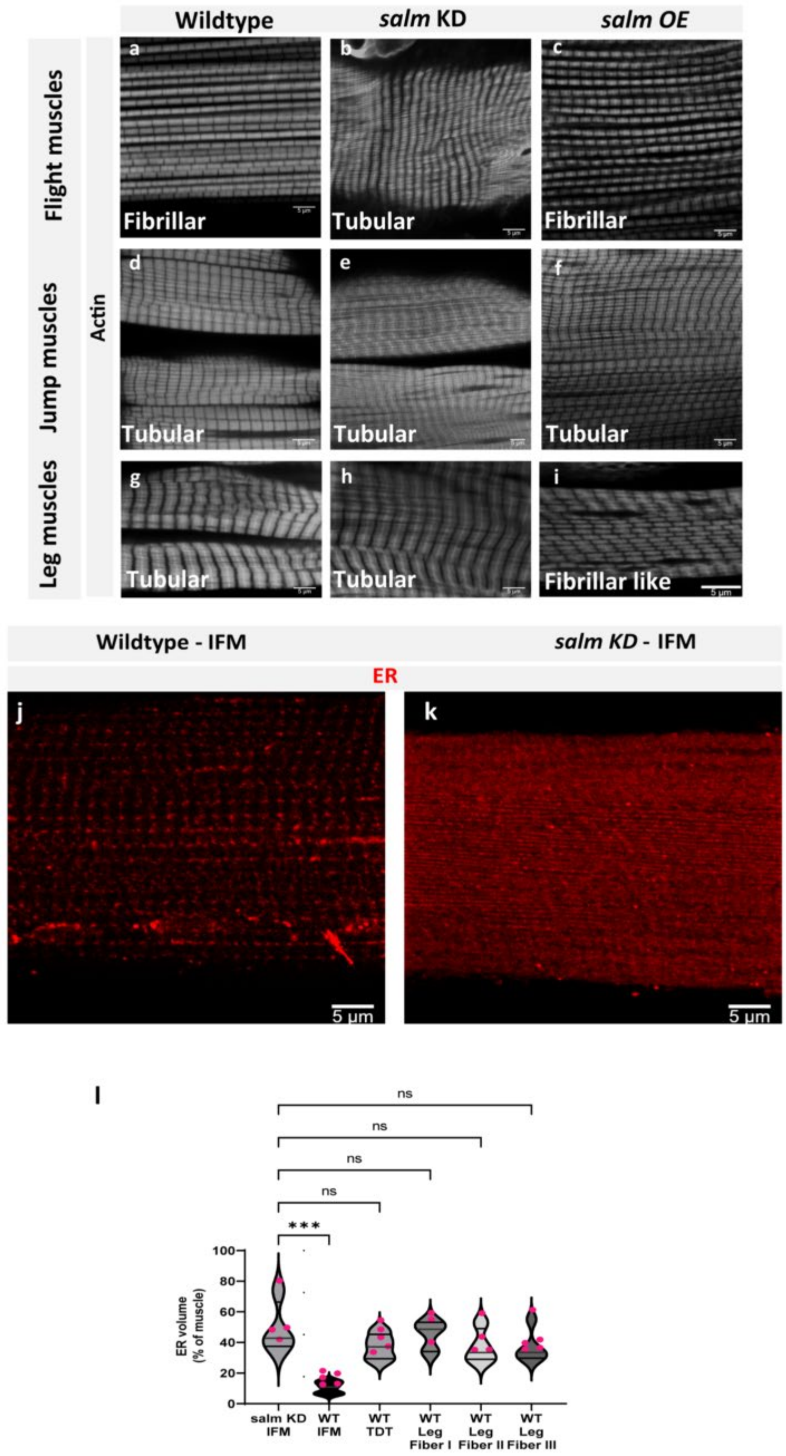


**Supplementary Fig. S8. *Salm* overexpression results in conversion of contractile type in leg muscles.** (a, b, c) Adult flight muscles (IFM) stained for F-actin (phTRITC) showing change in fiber type from (a) wildtype fibrillar to tubular fibers in (b) *salm* KD, but not in (c) *salm* OE. (d, e, f) Tubular jump muscles do not show alteration with either *salm* KD or *salm* OE. (g, h, i) Muscle contractile fiber type of wildtype leg muscles undergo conversion upon *salm* OE, but not *salm* KD. (j, k) Flight muscles stained for endoplamic reticulum (KDEL-rfp), mitochondria (mito-gfp), and nuclei (DAPI) showing increased ER content in (k) salm KD tubular IFMs compared to (j) wildtype fibrillar flight muscles (Scale bars: 5 µm). (l) Quantification of ER volume as a percent of muscle volume (*salm* KD-IFM, *n=3;* WT-IFM, *n=5*; Jump muscles (TDT), *n=5*; WT Leg Fiber I, *n=3*; WT Leg Fiber II, *n=4*; WT Leg Fiber III, *n=5*). Each point represents value for each dataset. Bars represent mean ± SD. Significance determined as $p < 0.05$ from one way ANOVA with Tukey's (*, $p \leq 0.05$; **,$p \leq 0.01$; ***,$p \leq 0.001$; ****, $p \leq 0.0001$; ns, non-significant)

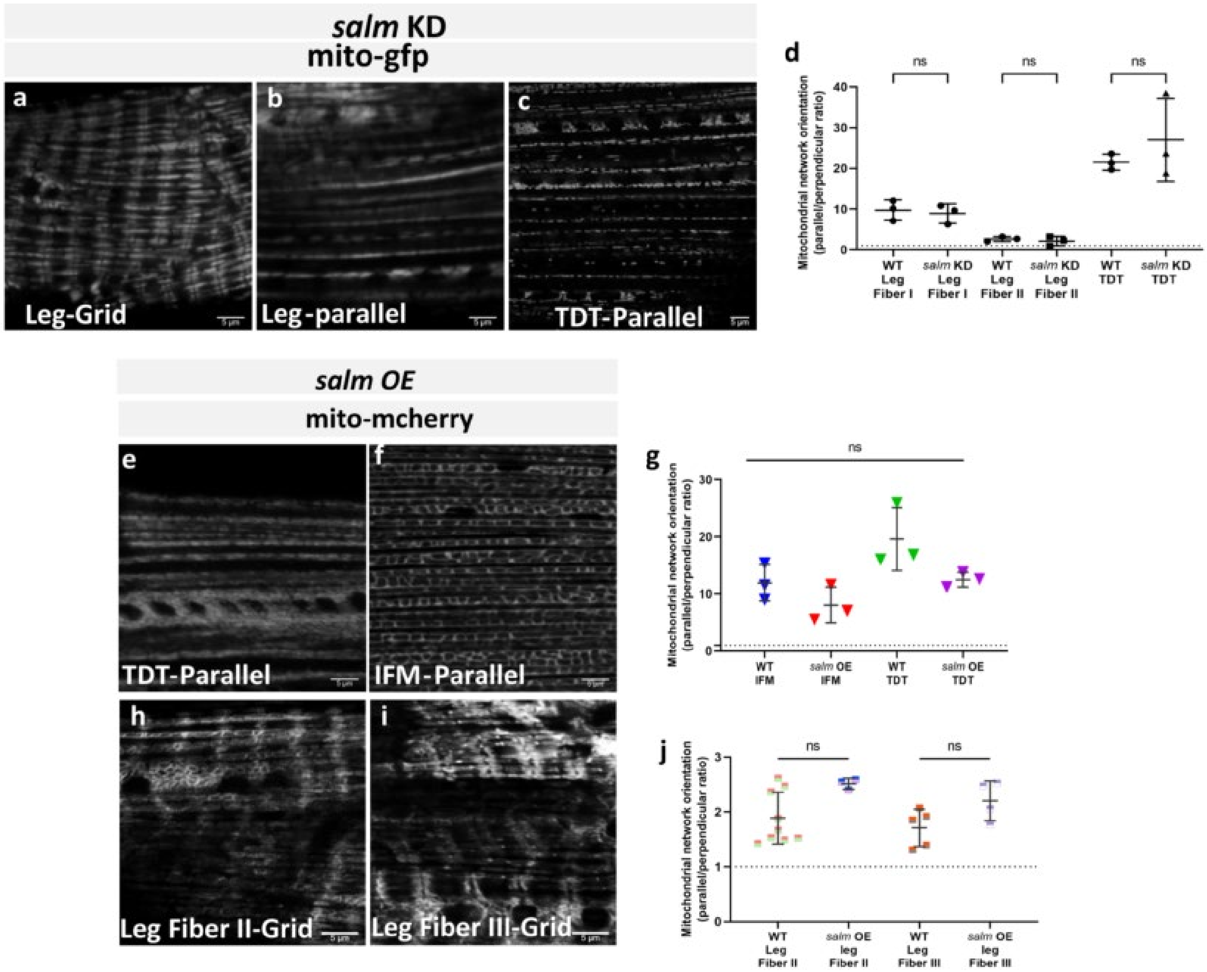


**Supplementary Fig. S9. Effect of s*alm* knockdown and overexpression on mitochondrial networks in muscles**. (a, b, c) Mitochondrial networks (mito-gfp) in *salm* KD (a, b) leg muscles and (c) jump (TDT) muscles (Scale bars: 5 µm). (d) Quantification of mitochondrial network orientation. Dotted line represents parallel equal to perpendicular. *mito-gfp;mito-mcherry;Mef2-Gal4* used as Wildtype (*n=3* for all groups). (e, f) Mitochondrial networks (mito-mcherry) in (e) jump and (f) flight muscles after *salm* OE (Scale bars: 5 µm for all). (g) Quantification of mitochondrial network orientation. Dotted line represents parallel equal to perpendicular. *mito-gfp;mito-mcherry;Mef2-Gal4* used as Wildtype (*n=3* for all groups). (h, i) Mitochondrial networks (mito-mcherry) in (h) Fiber II and (i) Fiber III of leg muscles after *salm* OE. (j) Quantification of mitochondrial network orientation in leg muscle fibers. Dotted line represents parallel equal to perpendicular. *mito-gfp;mito-mcherry;Mef2-Gal4* used as Wildtype (WT-Leg Fiber II, *n=9*; *salm* OE-Leg Fiber II, *n=3*; WT-Leg Fiber III, *n=5*; *salm* OE-Leg Fiber III, *n=4*). Each point represents value for each dataset. Bars represent mean ± SD. Significance determined as $p < 0.05$ from one way ANOVA with Tukey's (*, $p \leq 0.05$; **,$p \leq 0.01$; ***,$p \leq 0.001$; ****, $p \leq 0.0001$; ns, non-significant).

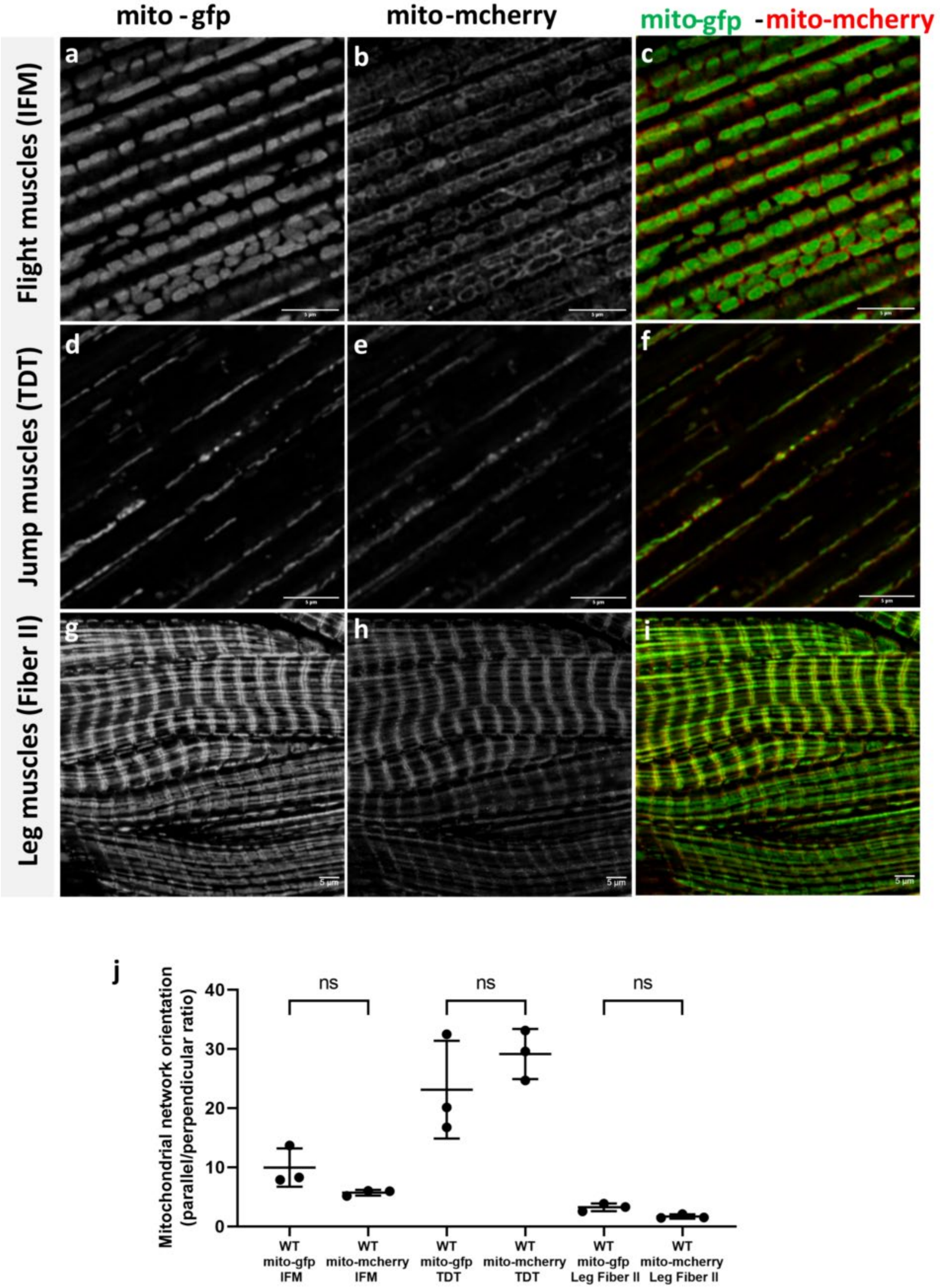


**Supplementary Fig. S10. Mitochondrial network organization in adult Drosophila muscles.** (a, b, c) Wildtype IFM with parallel mitochondria (mito-gfp and mito-mcherry). (d, e, f) Wildtype jump muscles showing parallel mitochondria. (g, h, i) Wildtype leg muscles showing grid-like mitochondria (mito-gfp and mito-mcherry) in Fiber II (Scale Bars: 5 µm). (j) Quantification of mitochondrial network orientation visualized by mito-gfp and mito-mcherry. Dotted line represents parallel equal to perpendicular. *mito-gfp;mito-mcherry;Mef2-Gal4* used as Wildtype (*n=3* for all groups) Bars represent mean ± SD. Significance determined as $p < 0.05$ from one way ANOVA with Tukey's (ns, non-significant).

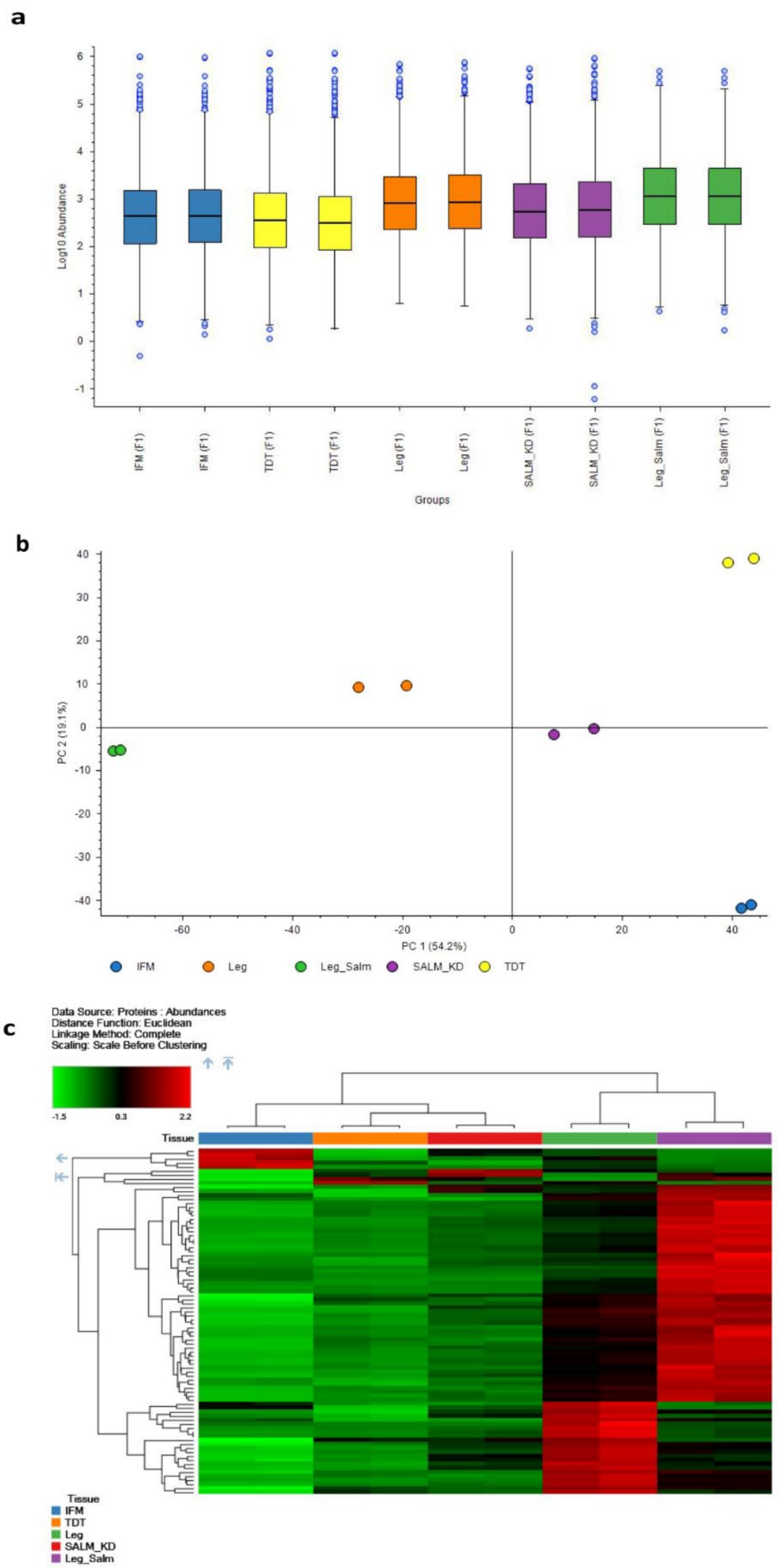


**Supplementary Fig. S11. Principal component analysis (PCA) and heatmap of proteome data.** (a) Comparison of protein abundance between muscle samples. The box-and-whisker plot shows the abundance of the intensity values for each individual muscle group. (b) Principal component analysis of the proteome data in a 2D graph of PC1 and PC2. (c) Heatmap of protein abundance pattern in differentially expressed proteins.

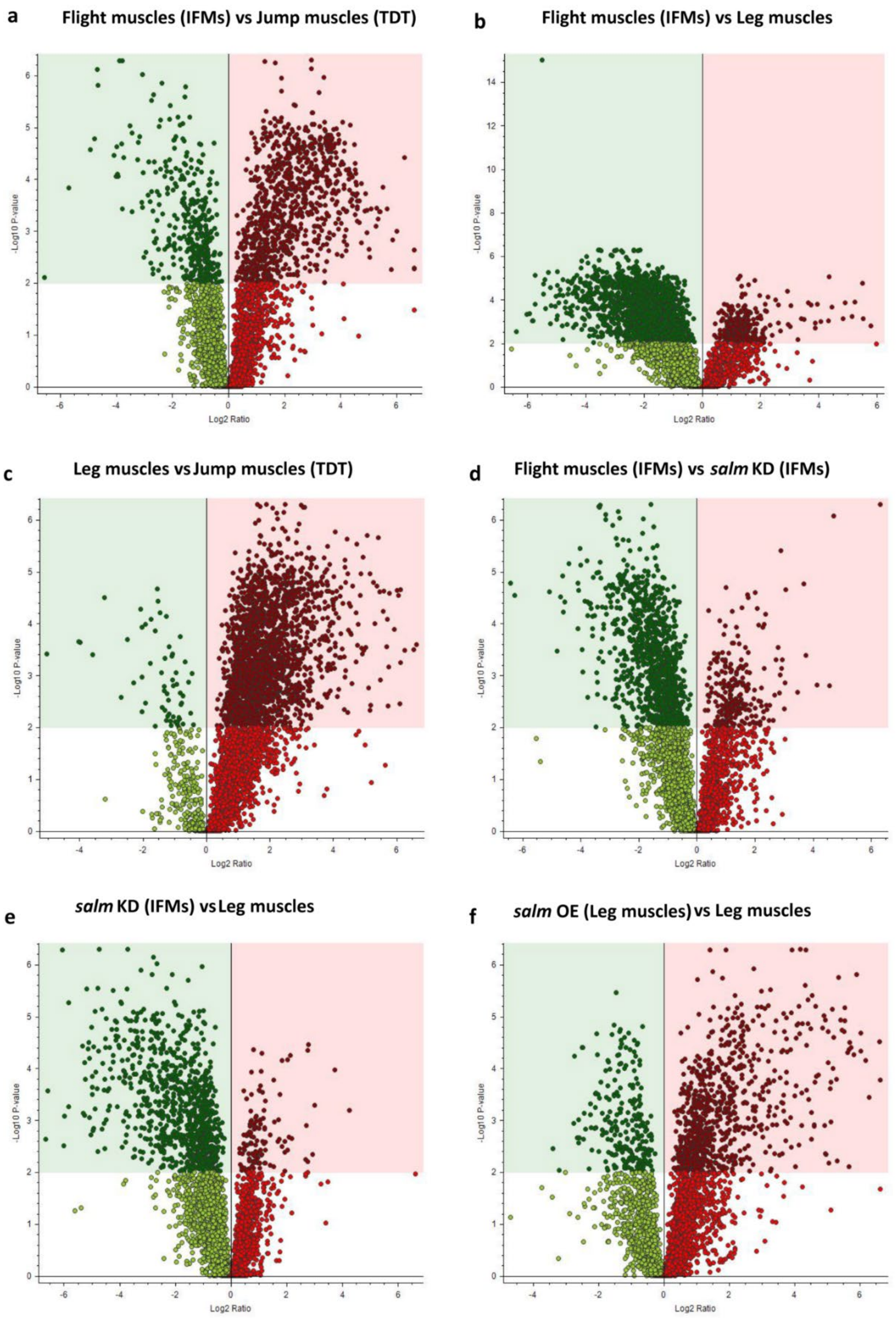


**Supplementary Fig. S12. Volcano plots showing differential protein expression between muscle types.** (a) Flight muscles (IFM) vs. jump muscles (TDT). (b) Flight muscles vs. leg muscles. (c) Leg muscles vs jump muscles. (d) Wildtype flight muscles vs. *salm* KD flight muscles. (e) *salm* KD flight muscles vs. wildtype leg muscles. (f) *salm* OE leg muscles vs. wildtype leg muscles. Green dots represent proteins with significantly lower abundances, while the red dots show proteins with higher levels of expression. X-axis, log 2 fold change (FC) differences in the gene expression and Y-axis, negative log 10 of the p values.

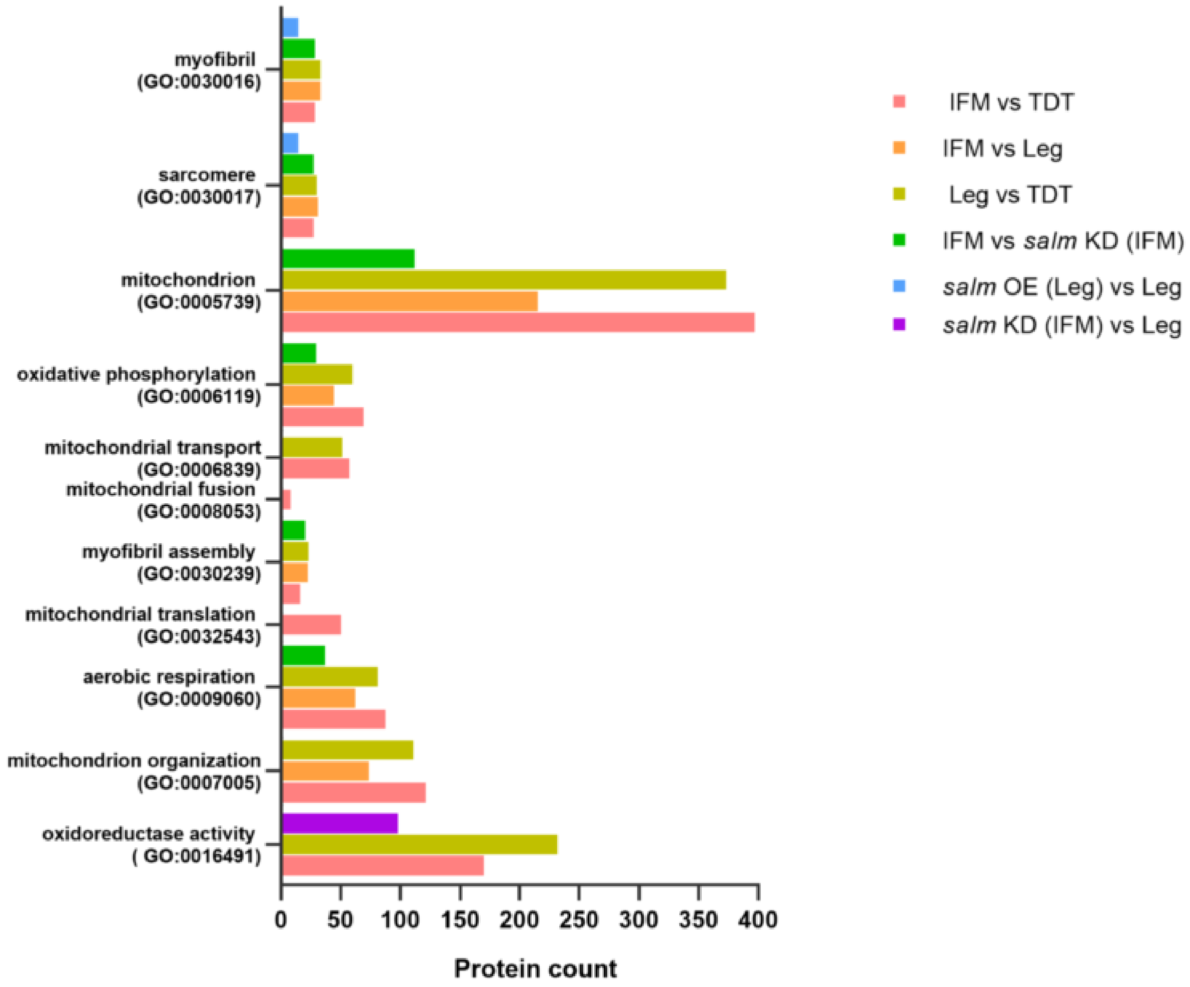


**Supplementary Fig. S13. Gene enrichment analysis of differentially expressed proteins among the five different muscle types using g:profiler.**

### Separating Contractile and Mitochondrial Fiber Type

| | | Flight | Jump | Leg | *salm KD*-Flight | *salm OE*-Leg |
|---|---|---|---|---|---|---|
| Contractile type | Fibrillar | + | - | - | - | + |
| | Tubular | - | + | + | + | - |
| Mito Network type | Parallel | ++ | ++ | + | - | + |
| | Grid | - | - | + | + | + |
| | Salm | ++ | + | - | - | + |

**Supplementary Fig. S14. Combinations of muscle contractile type, mitochondrial network configuration and *salm* expression among the five different muscle types.**

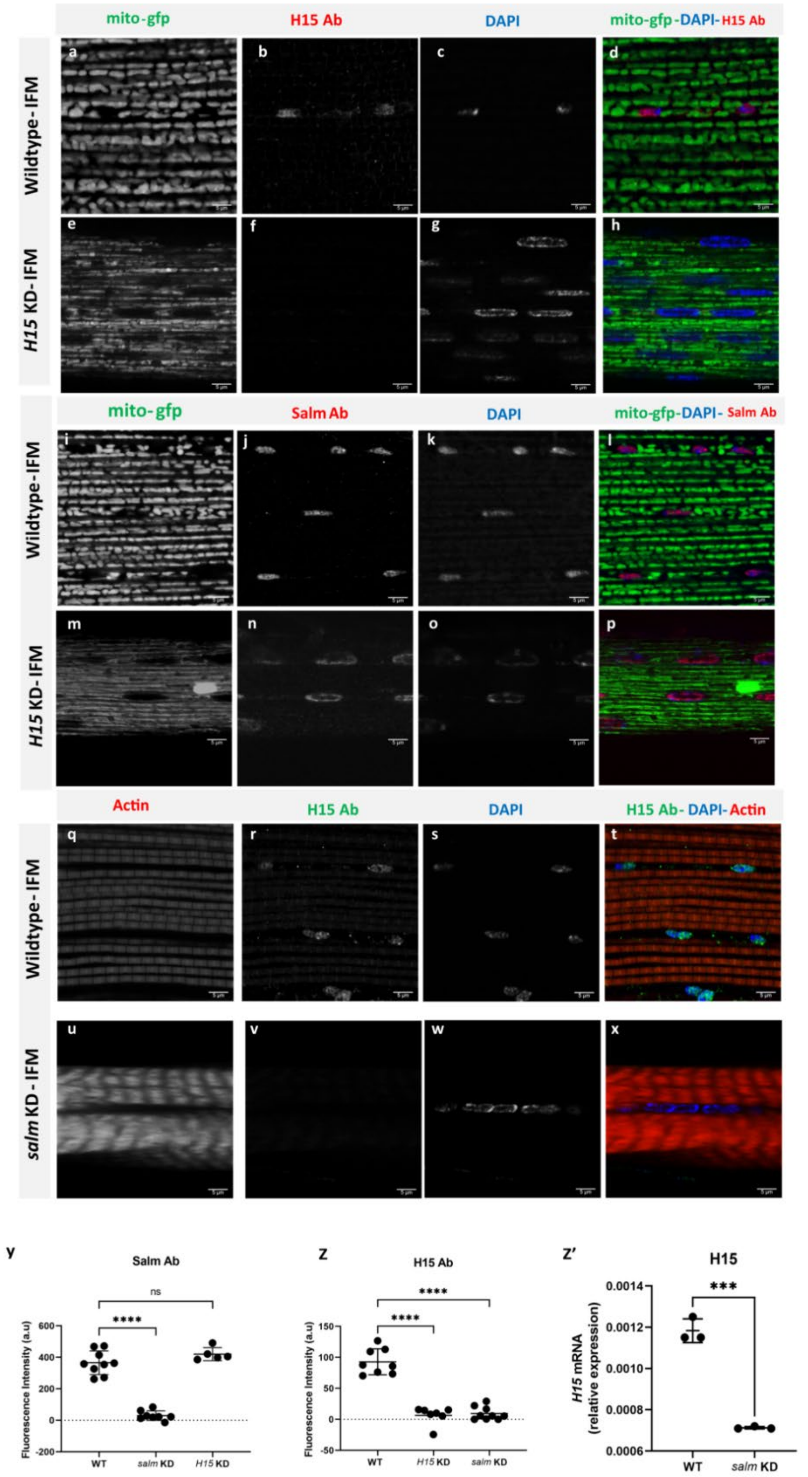


**Supplementary Fig. S15. *H15* is downstream of *salm* in the fiber type specification pathway.** (a-d) Wildtype IFM stained for mitochondria (mito-gfp), H15 antibody, and nuclei (DAPI) showing H15 expression in the nuclei. (e-h) *H15* KD IFM showing decreased expression of H15. (i-l) Wildtype IFM stained for mitochondria (mito-gfp), Salm antibody, and nuclei (DAPI) showing Salm expression in the nuclei. (m-p) In *H15* KD IFM, Salm expression is unaffected. (q-t) Wildtype IFM stained for mitochondria (mito-gfp), H15 antibody, and nuclei (DAPI). (u-x) *salm* KD IFM showing decreased expression of H15 (Scale Bars: 5 µm for all). (y, z) Quantification of fluorescence intensity of (y) Salm antibody staining and (z) H15 antibody staining. Each point represents value for each dataset. Bars represent mean ± SD. Significance determined as $p < 0.05$ from one way ANOVA with Tukey's (*, $p \leq 0.05$; **,$p \leq 0.01$; ***,$p \leq 0.001$; ****, $p \leq 0.0001$; ns, non-significant).

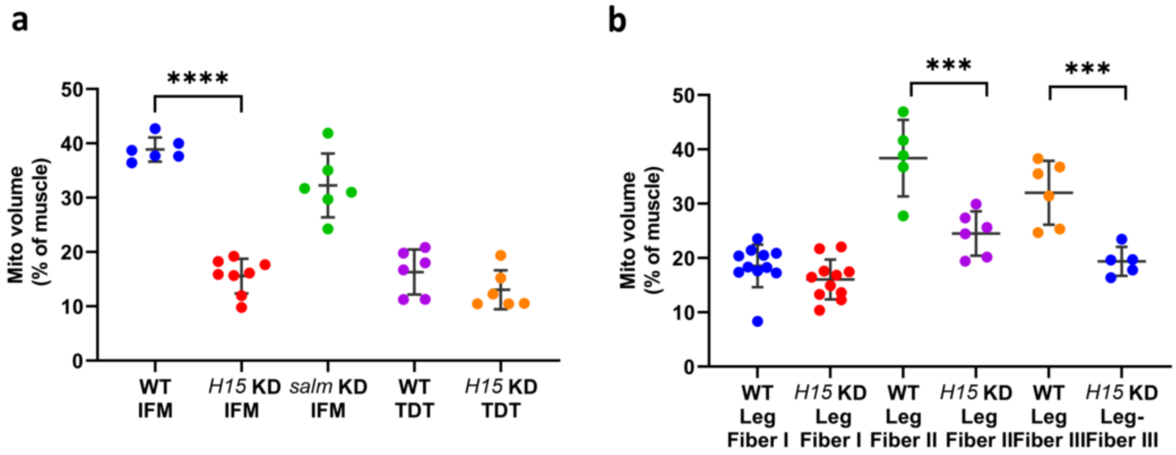


**Supplementary Fig. S16. Effect of H15 knock down on mitochondrial volume.** (a, b) Quantification of mitochondrial volume as a percent of total muscle volume in (a) IFM and (b) leg muscle fibers. *UAS-mito-gfp;UAS-mito-OMM-mcherry;Dmef2-Gal4* used as wildtype. (WT-IFM, *n=6*; *H15* KD-IFM, *n=7*; *salm* KD-IFM, *n=6*; WT-TDT, *n=6*; *H15* KD-TDT, *n=5*; WT-Leg Fiber I, *n=11*; *H15* KD-Leg Fiber I, *n=11*; WT-Leg Fiber II, *n=5*; *H15* KD-Leg Fiber II, *n=6;* WT-Leg Fiber III, *n=6*; *H15* KD-Leg Fiber III, *n=5*).

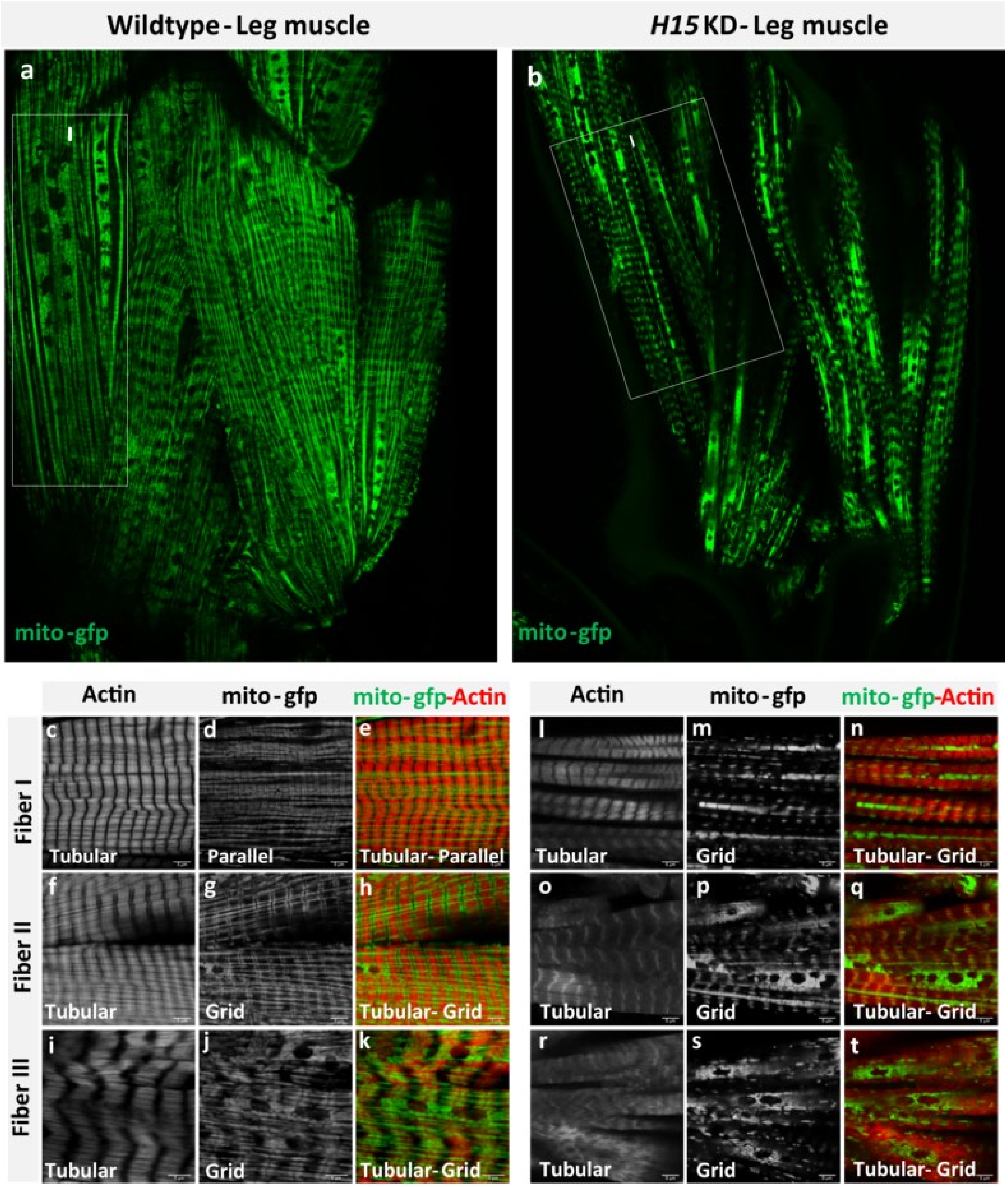


**Supplementary Fig. S17. *H15* regulates conversion of mitochondrial networks, but not contractile type in Drosophila leg muscles.** (a) Adult wildtype leg coxa muscles showing parallel mitochondrial networks (mito-gfp) in Fiber I (marked within rectangle) and grid-like networks in other regions. (b) *H15* KD leg muscles showing a uniform grid-like mitochondrial networks, even in fiber I (marked by rectangle) (Scale bar: 20 µm). (c, d, e) Wildtype leg muscle Fiber I showing parallel mitochondrial networks (mito-gfp) and tubular muscle fiber (phTRITC). (f, g, h) Wildtype leg muscle Fiber II and (i, j, k) Fiber III showing grid-like mitochondrial networks and tubular myofibrils. (l, m, n) *H15* KD leg muscles Fiber I showing grid-like mitochondrial networks, unlike wildtype Fiber I, but myofibrils remain tubular. (o, p, q) H15 KD leg muscle Fiber II and (r, s, t) Fiber III showing grid-like mitochondrial networks and tubular muscle fibers as in wildtype (Scale Bars: 5 µm).

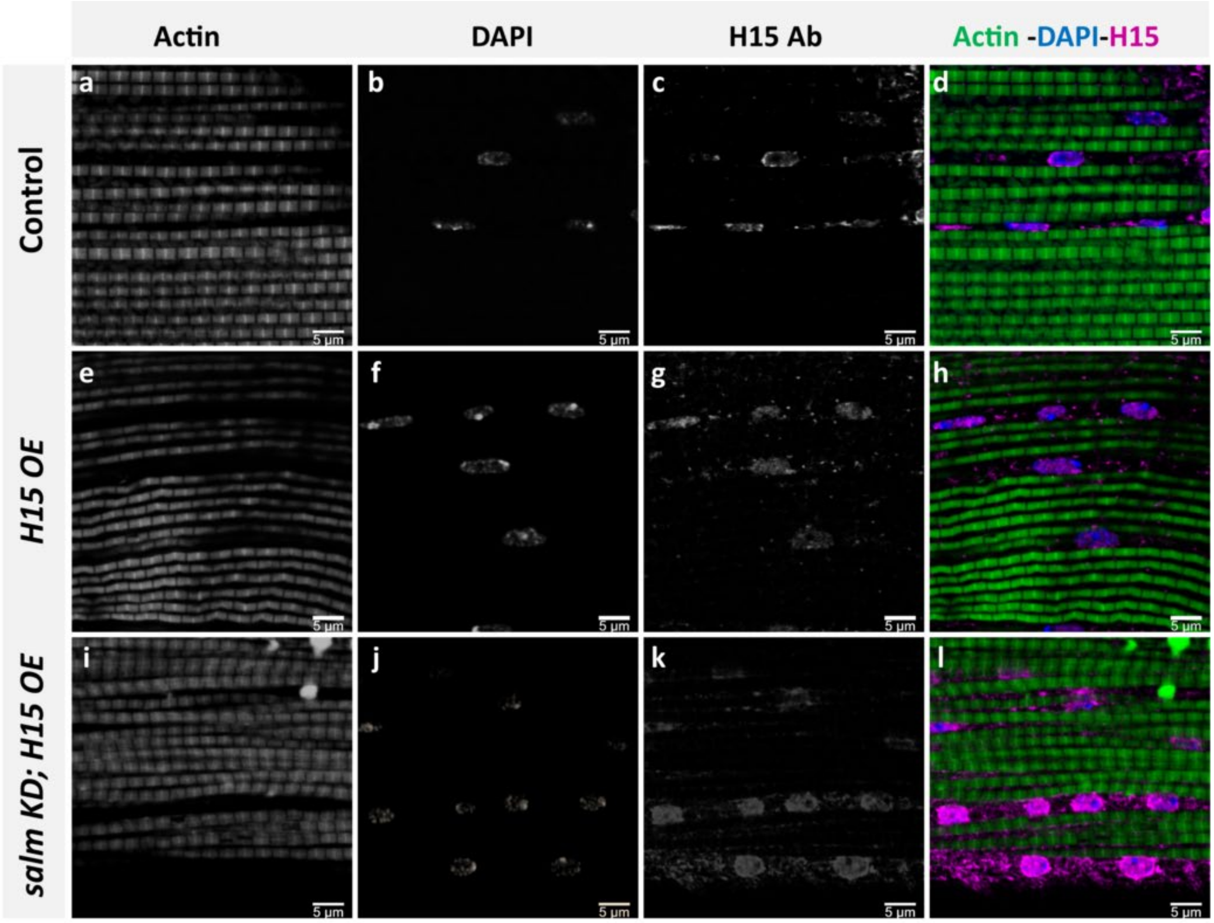


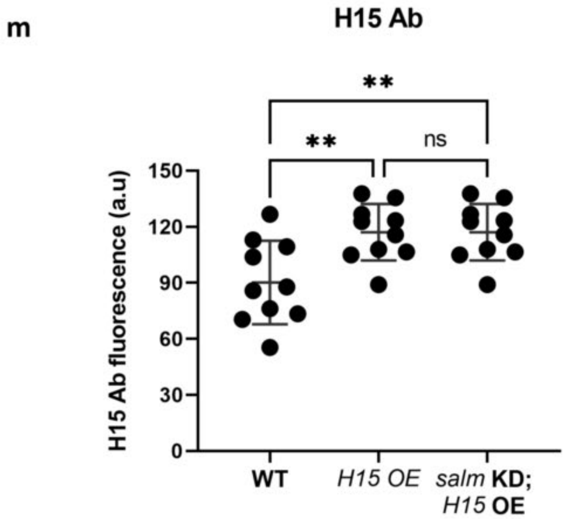


**Supplementary Fig. S18. *H15* overexpression in *salm* knock down background.** (a-d) Wildtype fibrillar flight muscles (IFMs) stained for F-actin (phTRITC), nuclei (DAPI), and H15 antibody showing H15 expression in the nuclei. (e-h) *H15* OE IFM showing increased expression of H15. (i-l) *H15* OE; *salm* KD IFM showing overexpression of H15 (Scale Bars: 5 µm). (m,n) Quantification of fluorescence intensity of (m) H15 antibody staining. Each point represents value for each dataset. Bars represent mean ± SD. Significance determined as $p < 0.05$ from one way ANOVA with Tukey's (*, $p \leq 0.05$; **,$p \leq 0.01$; ***,$p \leq 0.001$; ****, $p \leq 0.0001$; ns, non-significant).

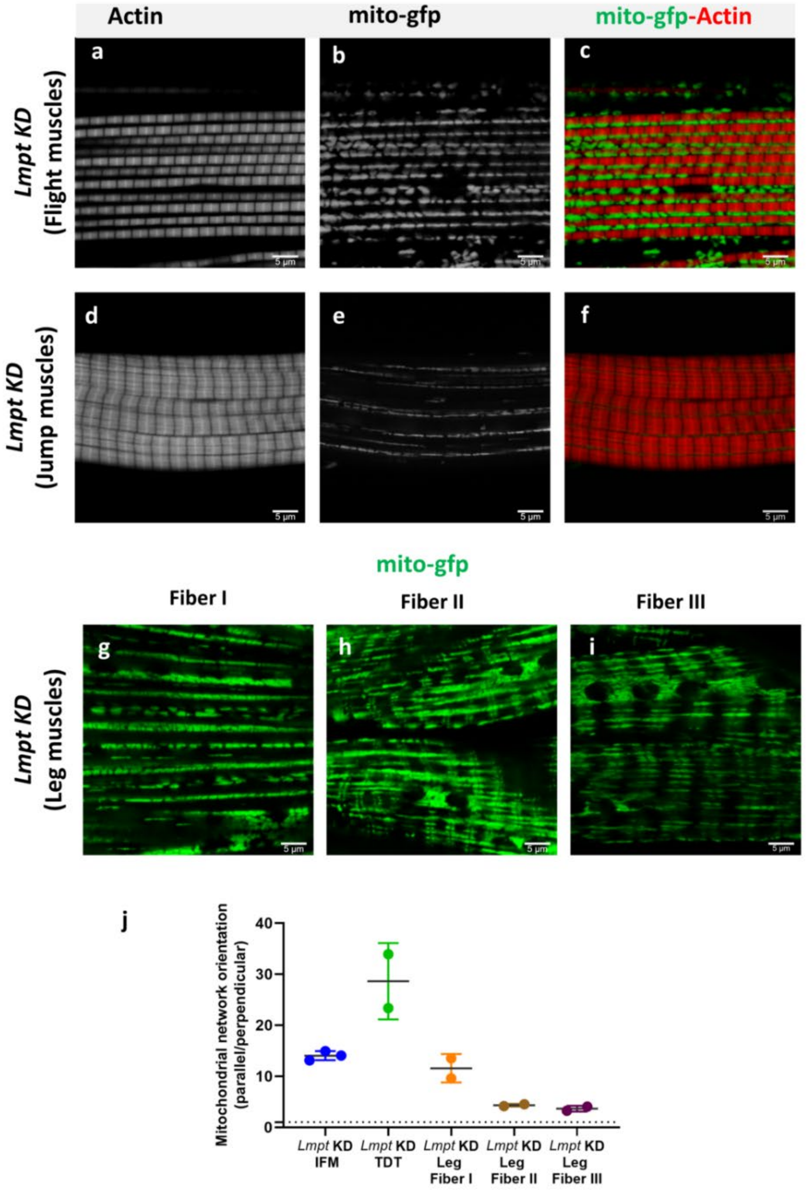


**Supplementary Fig. S19. *lmpt* knock down in muscles does not affect mitochondrial networks and contractile type in Drosophila muscles.** (a, b, c) Adult flight muscles with *lmpt* KD showing fibrillar myofibrils (ph-TRITC) and parallel mitochondrial networks (mito-gfp). (d, e, f) *lmpt* KD jump muscles showing tubular contractile type and grid-like mitochondrial network. (g) Leg muscle Fiber I with *lmpt* KD retains parallel mitochondrial networks (mito-gfp). (h) *lmpt* KD leg muscle Fiber II and (i) *lmpt* KD Fiber III showing grid-like mitochondrial networks (Scale Bars: 5 µm). (j) Quantification of mitochondrial network orientation. Dotted line represents parallel equal to perpendicular.

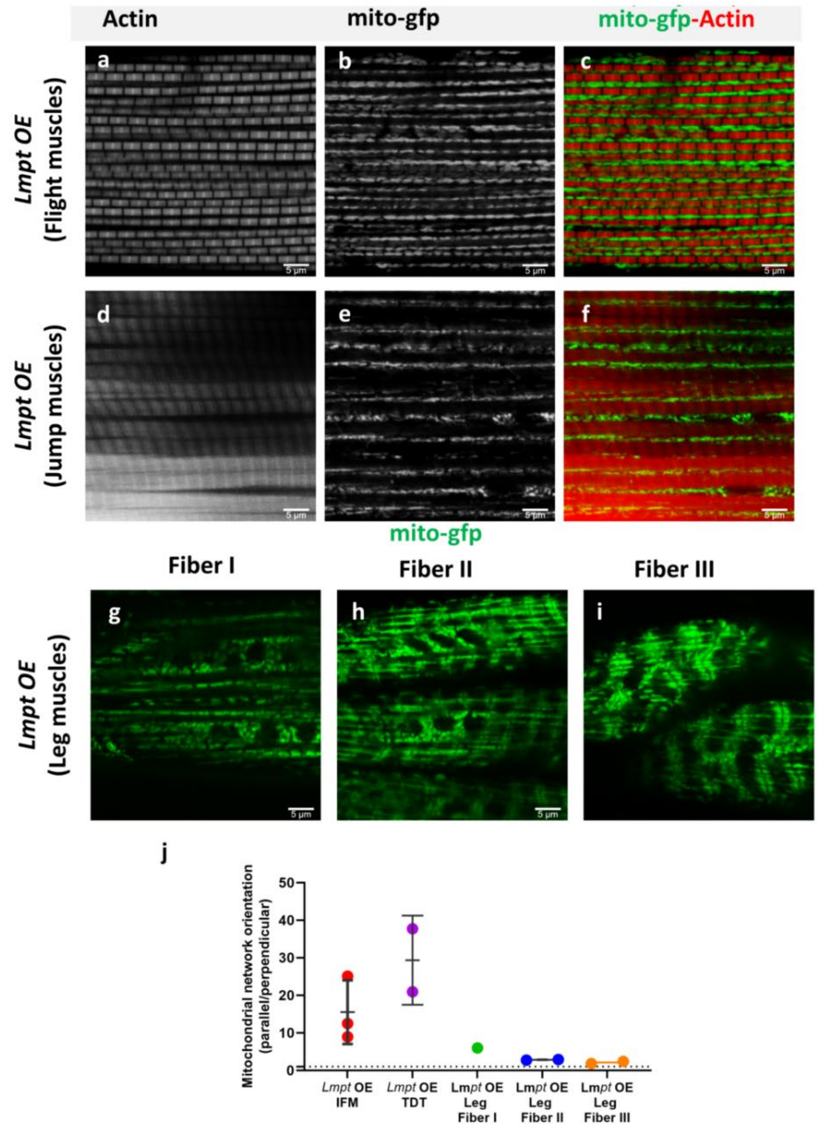


**Supplementary Fig. 20.** ***lmpt*** **overexpression in muscles does not affect mitochondrial networks and contractile type in Drosophila muscles.** (a, b, c) Adult flight muscles with *lmpt* OE showing fibrillar myofibrils (ph-TRITC) and parallel mitochondrial networks (mito-gfp). (d, e, f) *lmpt* OE jump muscles showing tubular contractile type and grid-like mitochondrial network. (g) Leg muscle Fiber I with *lmpt* OE retains parallel mitochondrial networks (mito-gfp). (h) *lmpt* OE leg muscle Fiber II and (i) *lmpt* OE Fiber III showing grid-like mitochondrial networks (Scale Bars: 5 µm). (j) Quantification of mitochondrial network orientation. Dotted line represents parallel equal to perpendicular.

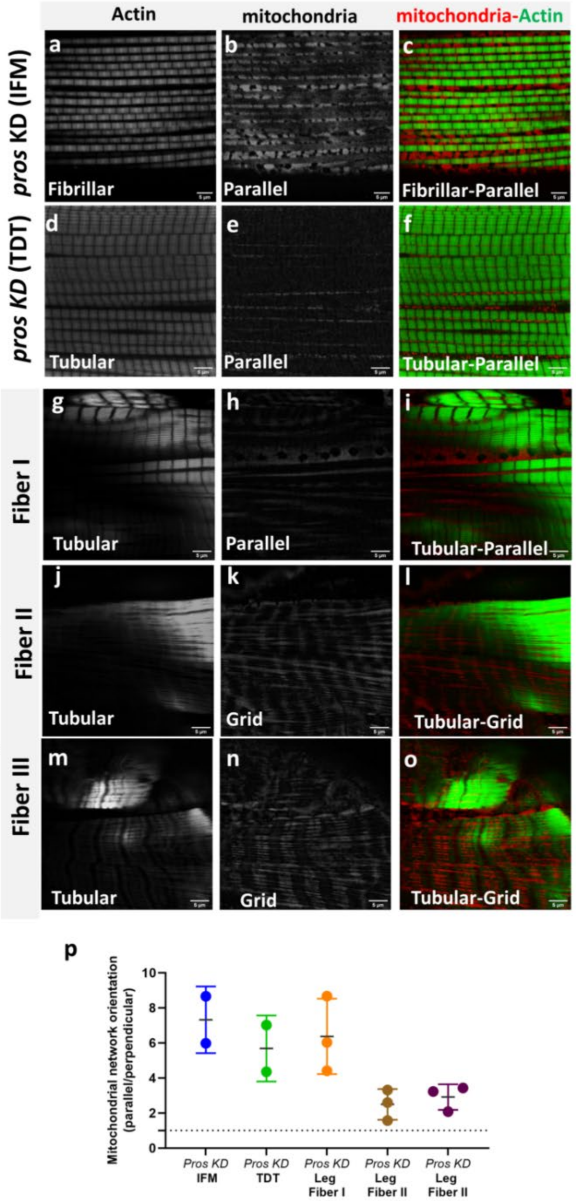


**Supplementary Fig. 21.** ***Pros*** **knock down in muscles does not affect mitochondrial networks and contractile type in Drosophila muscles.** (a, b, c) Adult flight muscles with *Pros* KD showing fibrillar myofibrils (ph-TRITC) and parallel mitochondrial networks (mito-gfp). (d, e, f) *Pros* KD jump muscles showing tubular contractile type and grid-like mitochondrial network. (g, h, i) Leg muscle Fiber I with *Pros* KD retains parallel mitochondrial networks and tubular muscle fibers. (j, k, l) *Pros* KD leg muscle Fiber II and (m, n, o) *Pros* KD Fiber III showing grid-like mitochondrial networks and tubular myofibrils (Scale Bars: 5 µm). (j) Quantification of mitochondrial network orientation. Dotted line represents parallel equal to perpendicular.

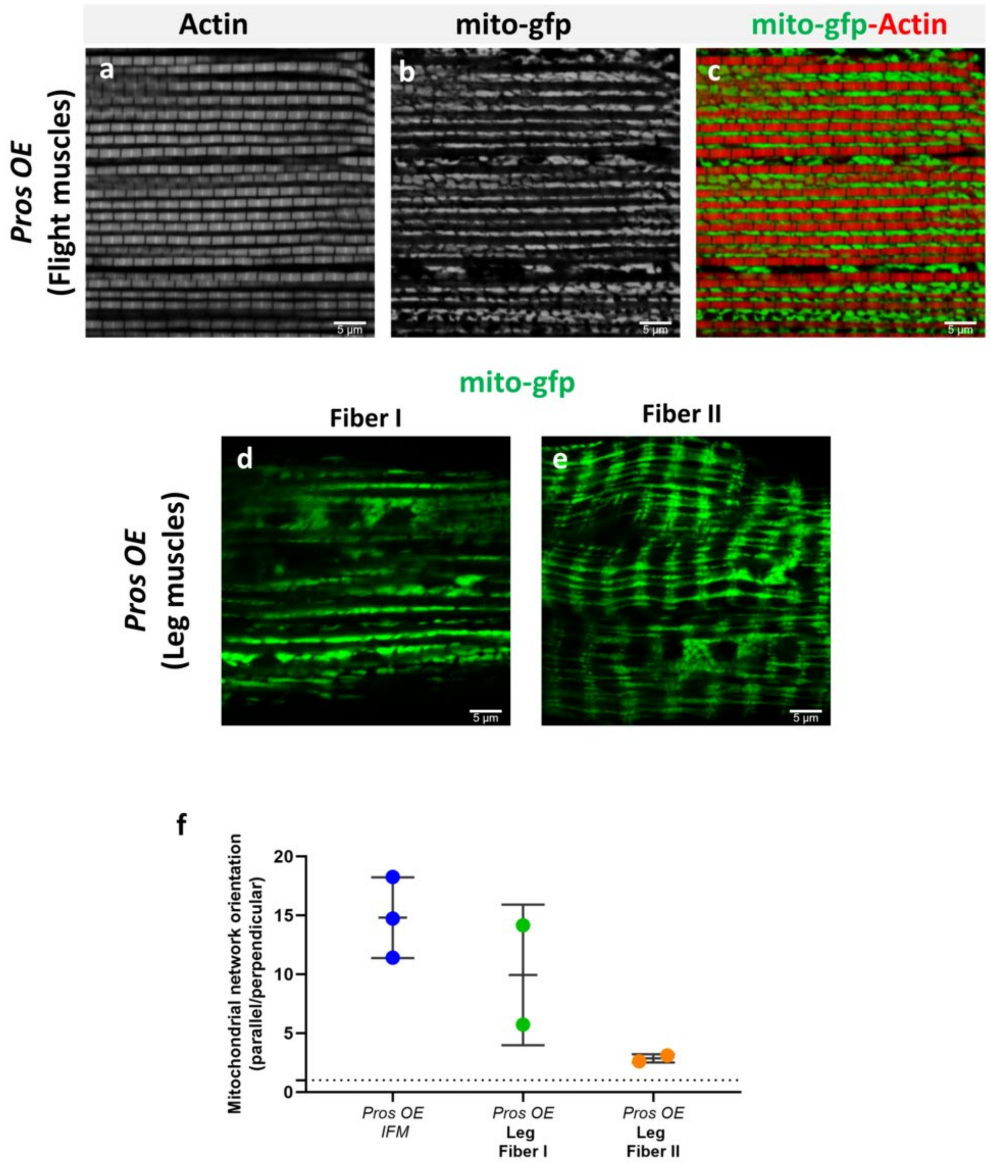


**Supplementary Fig. 22.** ***Pros*** **overexpression in muscles does not affect mitochondrial networks and contractile type in Drosophila muscles.** (a, b, c) Adult flight muscles with *Pros* OE showing fibrillar myofibrils (ph-TRITC) and parallel mitochondrial networks (mito-gfp). (d) Leg muscle Fiber I with *Pros* OE retains parallel mitochondrial networks (mito-gfp). (e) *Pros* OE leg muscle Fiber II showing grid-like mitochondrial networks (Scale Bars: 5 µm). (f) Quantification of mitochondrial network orientation. Dotted line represents parallel equal to perpendicular.

**WT Direct flight muscles**

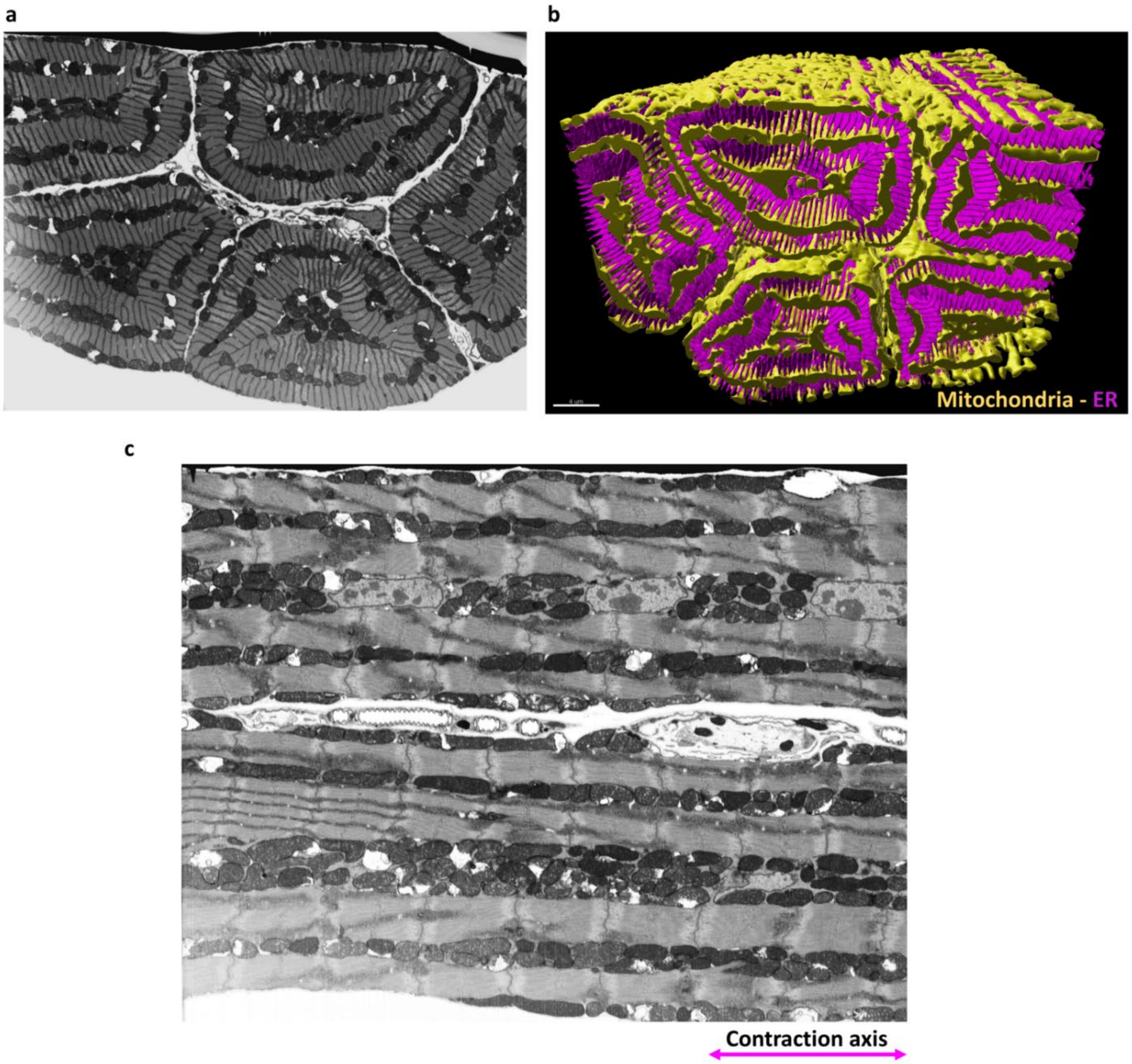


**Supplementary Fig. 23. Focused Ion bean scanning electron microscopy (FIB-SEM) images of Drosophila direct flight muscles.** (a) Transverse view of mitochondrial organization in wildtype direct flight muscles. (b) Representative 3D rendering of electron microscopic images of mitochondrial arrangement (yellow) and ER (magenta) in wildtype direct flight muscles showing mitochondrial networks arranged in sheets parallel to the tubular contractile networks. (c) Longitudinal view of mitochondrial organization in wildtype direct flight muscles.

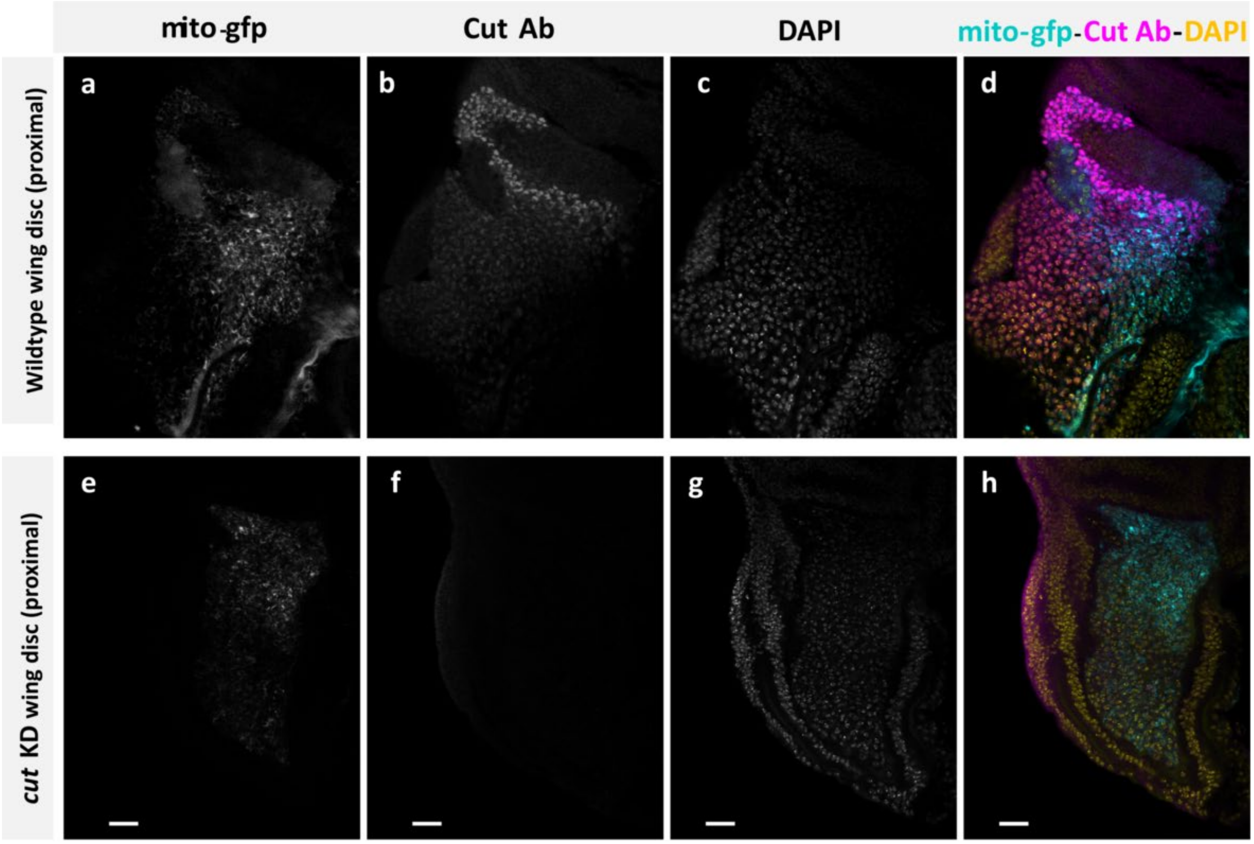


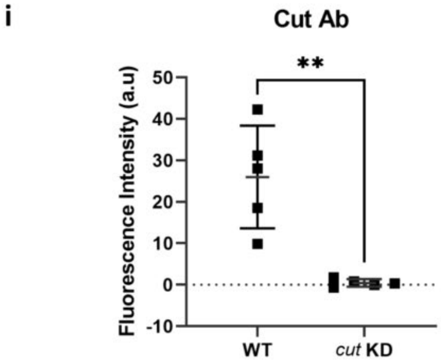


**Supplementary Fig. S24. Cut expression in wildtype and *cut* knock down wing imaginal discs.** (a-d) Wildtype wing imaginal discs stained for mitochondria (mito-gfp), Cut antibody, and nuclei (DAPI). (e-h) *cut* KD wing imaginal disc showing decreased Cut expression. (i) Quantification of fluorescence intensity of Cut antibody staining. Each point represents value for each dataset. Bars represent mean ± SD. Significance determined as $p < 0.05$ from unpaired t-test with Welch's correction (*, $p \le 0.05$; **,$p \le 0.01$; ***,$p \le 0.001$; ****, $p \le 0.0001$; ns, non-significant).

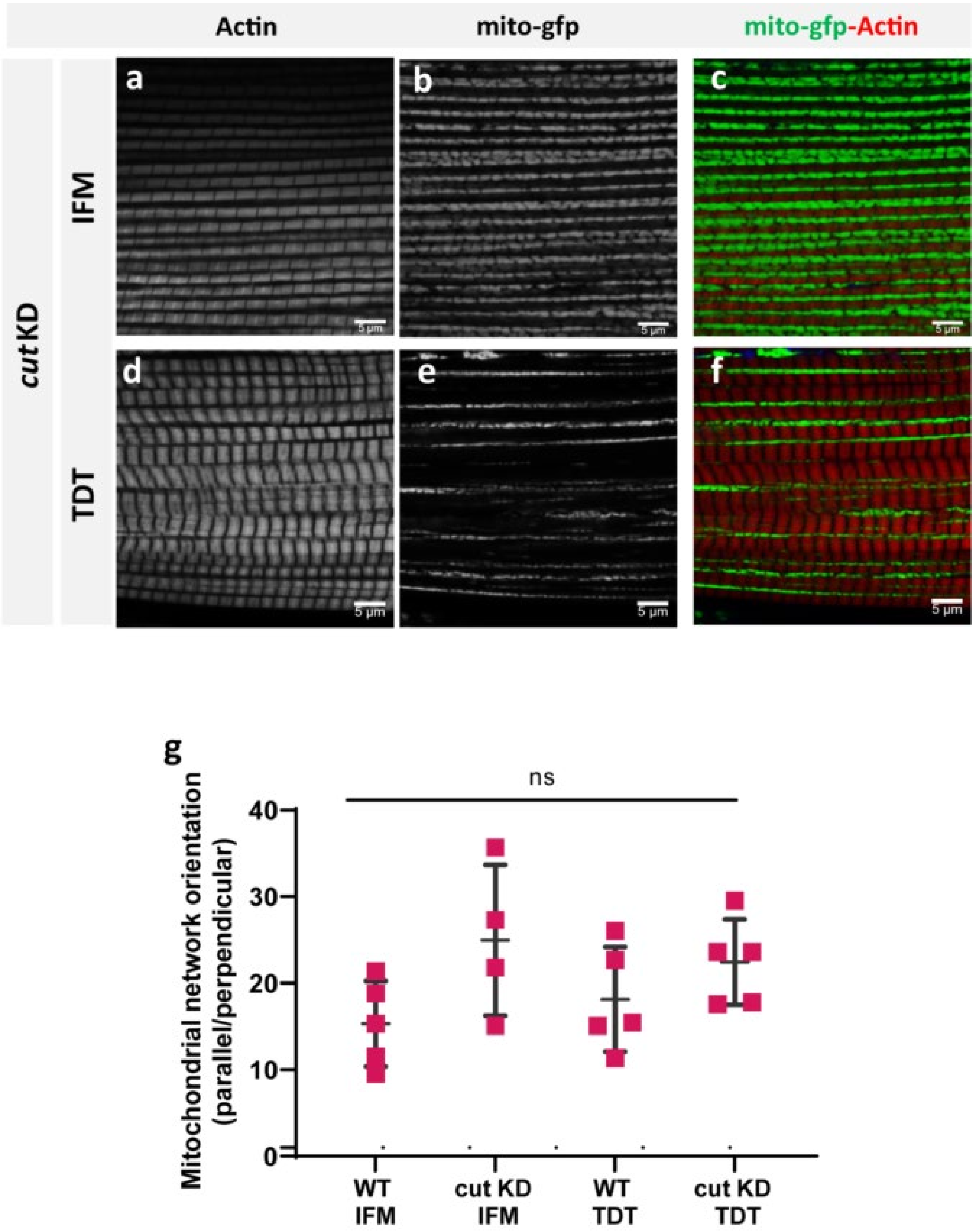


**Supplementary Fig. S25. *cut* knock down does not affect flight muscles and jump muscles.** (a, b, c) *cut* KD flight muscles (IFMs) showing parallel aligned mitochondria (mito-gfp) that are large, tube-like, and packed between fibrillar myofibrils (phTRITC). (d, e, f) *cut* KD jump muscles show mitochondria that are thin and elongated arranged in parallel mitochondrial networks between tubular fibers (Scale Bars: 5 µm). (g) Quantification of mitochondrial network orientation. Dotted line represents parallel equal to perpendicular (WT-IFM, *n=5*; *cut KD*-IFM, *n=4*; WT-TDT, *n=5*; *cut KD*- TDT, *n=5*). Each point represents value for each dataset. Bars represent mean ± SD. Significance determined as $p < 0.05$ from one way ANOVA with Tukey's (*, $p \leq 0.05$; **,$p \leq 0.01$; ***,$p \leq 0.001$; ****, $p \leq 0.0001$; ns, non-significant).

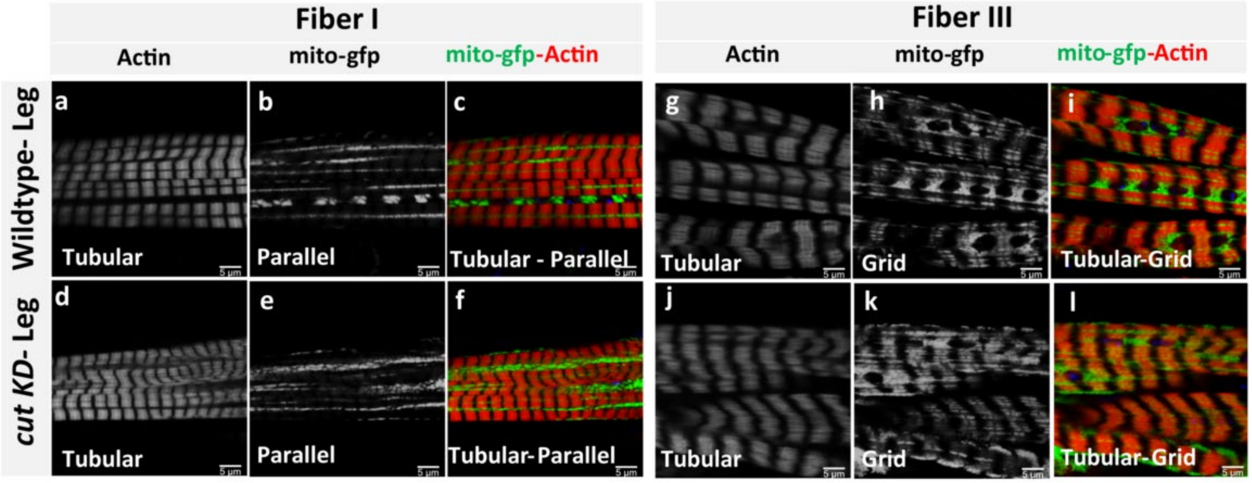


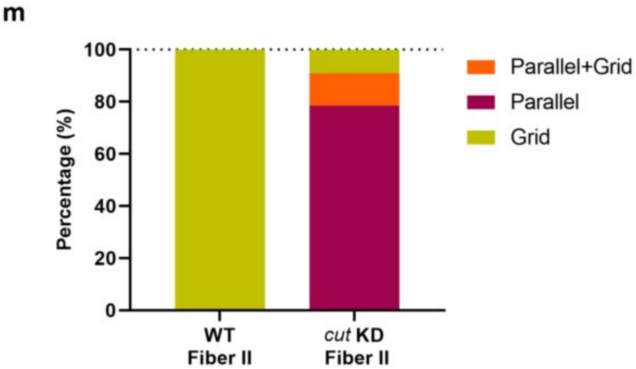


**Supplementary Fig. S26. *cut* does not regulate conversion of mitochondrial networks in Fibers I and III of Drosophila leg muscles.** (a, b, c) Fiber I of wildtype leg muscle showing parallel mitochondrial networks (mito-gfp) and tubular muscle fiber (phTRITC). (d, e, f) *cut* KD Fiber I of leg muscle showing parallel mitochondrial networks GFP) and tubular muscle fiber similar to wildtype Fiber I. (g, h, i) Fiber III of wildtype leg muscle showing grid-like mitochondrial networks and tubular muscle fiber. (j, k, l) *cut* KD Fiber III showing grid-like mitochondrial networks and tubular muscle fiber similar to wildtype Fiber III (Scale Bars: 5 µm). (m) Quantification of percentage of Fibers II exhibiting parallel, grid-like, and parallel as well as grid-like mitochondrial networks in wildtype and *cut* KD Fiber II of leg muscles (WT-Fiber II, *n=7*; *cut* KD-Fiber II, *n=14*).

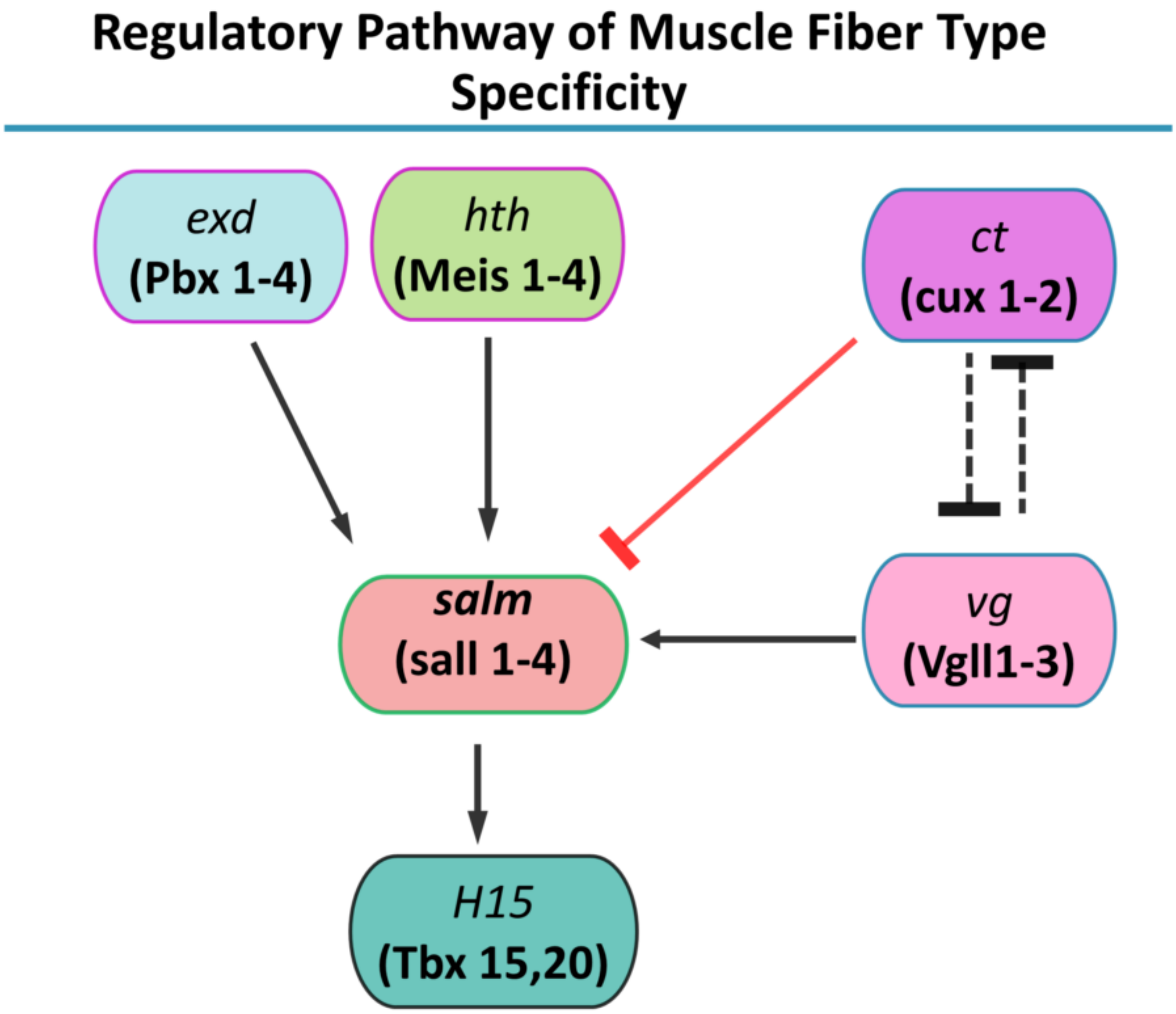


**Supplementary Fig. S27. Muscle fiber type specification pathway in Drosophila.** Evolutionarily conserved general regulatory pathway of muscle fiber type specificity in Drosophila. Parentheses show mammalian orthologs to Drosophila genes.
